# Life history adaptations to fluctuating environments: Combined effects of demographic buffering and lability of demographic parameters

**DOI:** 10.1101/2021.12.09.471917

**Authors:** Christie Le Coeur, Nigel G. Yoccoz, Roberto Salguero-Gómez, Yngvild Vindenes

## Abstract

Demographic buffering and lability have been identified as adaptive strategies to optimise fitness in a fluctuating environment. These are not mutually exclusive, however we lack efficient methods to measure their relative importance for a given life history. Here, we decompose the stochastic growth rate (fitness) into components arising from nonlinear responses and variance-covariance of demographic parameters to an environmental driver, which allows studying joint effects of buffering and lability. We apply this decomposition for 154 animal matrix population models under different scenarios, to explore how these main fitness components vary across life histories. Faster-living species appear more responsive to environmental fluctuations, either positively or negatively. They have the highest potential for strong adaptive demographic lability, while demographic buffering is a main strategy in slow-living species. Our decomposition provides a comprehensive framework to study how organisms adapt to variability through buffering and lability, and to predict species responses to climate change.

## Introduction

Understanding life history adaptations to fluctuating environments is increasingly important, as anthropogenic climate change is altering the temporal variability of multiple climatic drivers (IPCC, 2021; Laufkötter *et al*., 2020; Pendergrass *et al*., 2017). For instance, while an increased variance in daily and seasonal temperature and precipitation is expected across much of Europe in summer, a decrease is projected in other regions (Huntingford *et al*., 2013; IPCC, 2021; Kotz *et al*., 2021; Pendergrass *et al*., 2017). Fluctuations in abiotic and biotic environmental drivers experienced by organisms may affect their relative fitness and select for specific adaptations to live in variable environments.

Two main processes have been identified as adaptations to environmental variability, optimizing fitness: Demographic buffering reduces the variance in demographic parameters (e.g., survival, fertility), thereby minimizing the effects of bad environments (Morris & Doak, 2004; Hilde *et al*., 2020), while demographic lability lets the organisms take advantage of good environments by mounting a large increase in some demographic parameters compared to an average or bad environment, and therefore increasing their mean (Koons *et al*. 2009; Jongejans *et al*. 2010; Barraquand & Yoccoz 2013; see Box 1 for Glossary). The two processes are not mutually exclusive but can be selected simultaneously, so that different demographic parameters of a given life history can show different responses to an environmental driver. Yet, these processes have often been investigated separately, and we lack efficient methods to disentangle and predict their relative importance for a given life history and environment. To understand how organisms combine lability and buffering of their demographic parameters to enhance fitness in varying environments, we need a demographic model framework to predict two main fitness components: i) the effects of nonlinearity in responses of all demographic parameters to an environmental driver, and ii) the effects of variance-covariance of these parameters. While the latter is well described in stochastic demographic theory (Lande *et al*., 2003), we know much less about the impacts of nonlinearity, representing the potential for adaptation to varying environments through lability.

A key prediction from classical theory for evolutionary bet-hedging and stochastic population growth is that the long-term fitness will be reduced if the temporal variance of fitness is increased (Lewontin & Cohen, 1969). This result is assuming an unstructured population with annual population growth rates that are IID (independently and identically distributed). The fitness is then the logarithm of the geometric mean of these growth rates (Lewontin & Cohen, 1969). In structured populations, the stochastic growth process is more complex due to fluctuations in the (st)age structure that introduce autocorrelation in the annual growth rates (Caswell, 2001). Still, under the assumption of small fluctuations in the demographic parameters, Tuljapurkar (1982) derived an important approximation of long-term growth rate in stage-structured populations, emphasising how the variance in fitness is linked to variances and covariances of demographic parameters in different stages (equation 1). The key conclusion from this approximation is that temporal variability in demographic parameters and/or positive covariance will have a negative effect on fitness, and should be selected against, in particular for demographic parameters that have a large impact on fitness in the mean environment. Accordingly, the demographic buffering hypothesis predicts that natural selection should favour a reduction in variance of the demographic parameters with the strongest influence on population growth (Boyce *et al*., 2006; Gaillard & Yoccoz, 2003; Hilde *et al*., 2020; Pfister, 1998; Tuljapurkar & Orzack, 1980).

However, positive effects of environmental variability have also been demonstrated under strong negative covariance among demographic parameters (Colchero *et al*., 2019; Doak *et al*., 2005; Tuljapurkar, 1990), negative environmental autocorrelation (Tuljapurkar, 1982), and convex relationships between demographic parameters and the environment. The latter represents a case of adaptive lability as described by Koons *et al*. (2009). In contrast to adaptive demographic buffering, which optimizes fitness by reducing the variance of most influential demographic parameters, lability can be adaptive if the benefit of an increase in the arithmetic mean of the annual growth rates through increased demographic parameter means can overcome the negative effect of increased demographic variance on fitness (Box 1). Nonlinearity in population and demographic parameter responses to environmental drivers may be common in the wild (Barraquand & Yoccoz, 2013; Clark & Luis, 2020; Dahlgren *et al*., 2011; Drake, 2005; Hansen *et al*., 2021; Jenouvrier *et al*., 2012; Lawson *et al*., 2015; Louthan & Morris, 2021; Mysterud *et al*., 2001), highlighting the potential importance of lability as an adaptation to environmental variability. However, with structured life histories the combined effects of nonlinearity in different demographic parameters on fitness are challenging to predict (Koons *et al*., 2009).

Somewhat contrasting predictions have been made as to which demographic parameters should be labile or buffered, and the relative importance of each process for a given life history. Demographic lability has been suggested to affect mainly the demographic parameters with least effect on fitness (Hilde *et al*., 2020), as a consequence of selection for buffering of more influential demographic parameters. Other studies suggest that lability can be equally important to demographic buffering, and that it can also occur in highly influential demographic parameters (Jongejans *et al*., 2010; Koons *et al*., 2009; McDonald *et al*., 2017). Based on the latter prediction, recent research suggests that adaptive lability and buffering can be located at the opposite ends of a continuum, encompassing a wide range of demographic strategies (Salguero-Gómez, 2021; Santos *et al*., 2021). Yet, the extent to which lability among the least or the most influential demographic parameters can be adaptive strategies for coping with varying environments, relative to buffering, remains largely unexplored (e.g., Barraquand & Yoccoz 2013).

We thus need a more thorough understanding of how the opportunity for selection on demographic buffering and lability depends on major axes of life history variation such as the slow-fast continuum (Stearns, 1992; Gaillard *et al*., 2016; Salguero-Gómez *et al*., 2016b). For instance, populations of fast-living species have been predicted to be more responsive to environmental variability than those of slow-living species, and to be more likely to show adaptive lability in their demographic parameters (Dalgleish *et al*., 2010; Iles *et al*., 2019; Koons *et al*., 2009; Morris *et al*., 2008). According to demographic buffering hypothesis, species towards the slow end of the continuum benefit most from reduced variance in annual survival of the mature stages, while fast-living species gain relatively more from reduction of variance in annual fertility and/or survival of the immature stages (Hilde *et al*., 2020; Gaillard & Yoccoz, 2003; Rotella *et al*., 2012). These effects can be predicted from Tuljapurkar’s small noise approximation (Tuljapurkar, 1982; equation 1), but we lack a similar expression to describe the net impact of nonlinearity in different demographic parameters of the same life history. Here, we introduce a new ‘nonlinearity index’ to predict changes in the arithmetic mean arising from nonlinearity in different demographic parameter responses to an explicit environmental driver. We decompose the stochastic growth rate into contributions from nonlinearity effects and variancecovariance effects. We then apply the decomposition to study how organisms may combine adaptive buffering and lability responses depending on generation time, which closely correlates with the species’ position along the slow-fast continuum (Gaillard *et al*., 2005). We use population models from the COMADRE animal matrix database (Salguero-Gómez *et al*., 2016a) as a starting point for our calculations, representing a broad range of life histories in the mean environment. We then add stochastic environmental variation and perform the decomposition under different scenarios for nonlinearity and covariance among demographic parameters (Fig. 1). Our study provides a method to disentangle the effects of buffering and lability for any given life history, and the subsequent analysis addresses two main questions: First, what is the opportunity for positive effects due to adaptive lability to overcome negative impacts through the variance-covariance of demographic parameters, and how does this pattern depend on generation time? Second, are demographic parameters that show adaptive lability typically the least or most influential demographic parameters for fitness?

**Figure 1.**
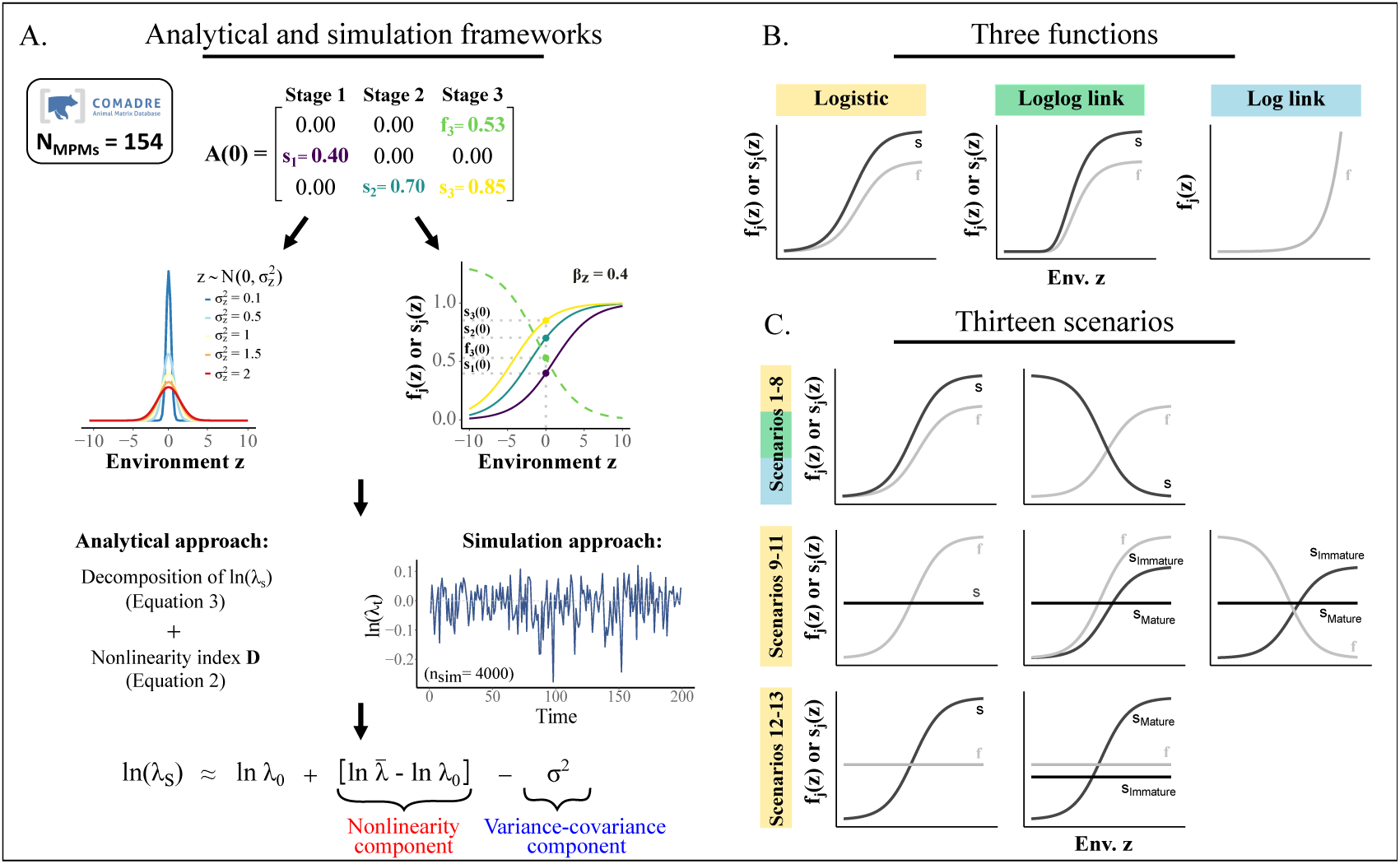
Framework used to study the effects of environmental variability on fitness (stochastic growth rate ln(*λ*_*s*_)). **A**. Our calculations define demographic parameters as nonlinear functions of the environmental driver *z* (see methods), where **A**(0) (from our selected, standardized COMADRE models, *N*_*tot*_= 154) defines the values of (st)age-specific survival rates *s*_*j*_(0) and fertilities *f*_*j*_(0) in the mean environment (z=0). Different levels of environmental variance levels 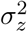 and environmental strength |*β*_*z*_| of *z* on the demographic parameters were considered. In the analytical approach, ln(*λ*_*s*_) was calculated and decomposed into main components capturing nonlinearity and variance-covariance effects following equation 3. The accuracy of this decomposition was tested using simulations (Supporting information S4). **B**. Two or three different link-functions were considered for survival *s*_*j*_(*z*) and fertility *f*_*j*_(*z*), respectively. **C. Scenarios 1-8:** Four combinations were examined including logistic functions for all parameters, loglog link functions for all parameters and two combinations of exponential fertilities *f*_*j*_(*z*) (log link) with logistic or loglog link function for *s*_*j*_(*z*). Positive or negative covariance between survival and fertility was tested for each combination, assuming positive covariance between *s*_*j*_(*z*), and between *f*_*j*_(*z*). **Scenarios 9-11:** Scenarios of forced buffering considering demographic lability in the fertility coefficients and survival rates of the immature stages (*S*_*immature*_). **Scenarios 12-13:** Scenarios of forced buffering assuming demographic lability in all survival rates *s*_*j*_(*z*) or in only the mature stages (*S*_*mature*_). Logistic functions were used to define lability while the other rates were held constant and fixed to the values reported in the standardized COMADRE projection matrix.

## Material and methods

To explore fitness responses to environmental variability along the slow-fast continuum, we decomposed the long-term stochastic growth rate ln(*λ*_*s*_), a measure of fitness (Tuljapurkar, 1990; Caswell, 2001; Lande *et al*., 2003), into main components capturing effects of nonlinearity in demographic parameters as a function of an environmental driver *z*, and effects of variance-covariance among the parameters. Our approach builds on Tuljapurkar’s approximation which assumes linear relationships between demographic parameters and an IID environmental variable: (Tuljapurkar, 1990):

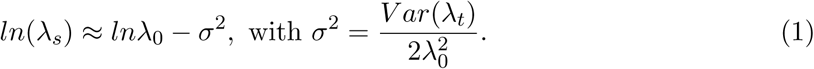

Here λ_0_ is the arithmetic growth rate in the mean environment, which is assumed equal to the mean arithmetic growth rate 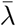 (ignoring non-linear responses), while *V ar*(λ_*t*_) is the variance in annual population growth caused by temporal variance and covariance in the demographic parameters. We show in the next section that including nonlinear effects of the environment on demographic parameters mainly affects ln(λ_*s*_) through the mean arithmetic growth rate 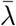, but also through the variance-covariance term. After defining main components of the stochastic growth rate, we perform a theoretical exploration of how the different components will vary across generation time, using different scenarios regarding nonlinear functions for survival and fertility (Fig. 1A-C). We also confront hypotheses about demographic lability, through scenarios that specifically consider effects of nonlinearity in the demographic parameters of immature or mature individuals only, keeping other parameters constant (’forced buffering’ scenarios, Fig. 1C). All simulations and calculations were performed in R, version 4.0.3 (R Core Team, 2020). R code is provided in Supporting information S7.

### Decomposing the stochastic growth rate with nonlinear effects

We assume that the environment at each time step is described by a stochastic variable *z* (IID), with mean 0 and variance 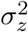. Population growth from one time step to the next is given by **n**_*t*+1_ = **A**(*z*_*t*_)**n**_*t*_, where **n**_*t*_ is the vector containing the number of individuals in each stage at time *t*, and **A**(*z*) is the population projection matrix. Elements of **A**(*z*) are the demographic parameters describing survival, fertility and transitions as functions of *z*. To derive the stochastic growth rate, we approximate this projection matrix using 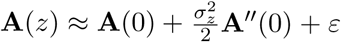, where *ε* is the matrix describing the noise terms with mean elements 0, **A**(0) is the projection matrix of the mean environment (with asymptotic growth rate λ_0_) and **A**″(0) = **A**″(*z*)|_*z*=0_ contains the second derivatives of elements of **A**(*z*). Using this second derivative matrix, we define a nonlinearity index (Supporting information S3)

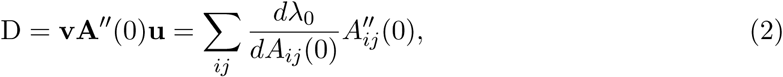

which measures the overall degree of nonlinearity in the life history defined by **A**(*z*). A positive *D* indicates adaptive lability. A matrix element (i.e., demographic parameter) with strong convex curvature may still have a low impact on *D* if the corresponding sensitivity of λ_0_ to that element is low, and vice versa.

Applying a Taylor approximation to the mean change of the logarithm of the total reproductive value *V*_*t*_ =∑ _*j*_ *n*_*j,t*_*v*_*j*_ (where reproductive values **v** are calculated for the matrix **A**(0)), we show in Supporting information S3 that the stochastic growth rate is given by

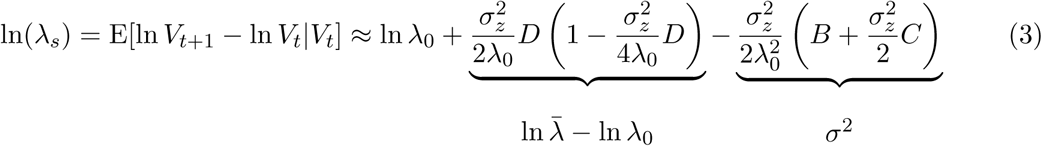

where *D* is the nonlinearity index defined above, 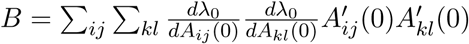 (where **A**′(0) = **A**′(*z*)|_*z*=0_ is the matrix of first derivatives), and 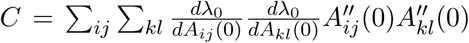. The stochastic growth rate is thus decomposed into the growth rate in the mean environment, ln λ_0_, plus two additive terms describing changes mainly due to nonlinearity 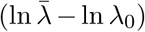, and changes mainly due to variance-covariance (*σ*^2^) of demographic parameters in a stochastic environment. The first term can be positive or negative, depending on the nonlinearity index *D*, and can be further decomposed into effects of survival and fertility coefficients (code in Supporting information S7). The second term corresponds largely to the variance-covariance term in the approximation of Tuljapurkar (1982), except that here there is also a small effect of nonlinearity through *C*. However, the effect of nonlinearity on the second term is very small compared to the effect of nonlinearity on the mean, therefore we refer to the first term as the nonlinearity component and second term as the variance-covariance component. In the Supporting information S4 we demonstrate the accuracy of this approximation using simulations.

### Applying the decomposition

To explore life history variation in the main components of the stochastic growth rate, we used age-and stage-structured matrix population models (MPMs) from the COMADRE Animal Matrix Database (v.4.20.5; Salguero-Gómez *et al*., 2016a) as a starting point, considering different scenarios for effects of the environment *z* on the demographic parameters. Each MPM includes a projection matrix that depends on the (st)age-specific fertilities, transitions, and survival rates for a given time interval (see Fig. 1). We let this projection matrix represent the matrix in the mean environment, **A**(0). We selected MPMs from unmanipulated and free-ranging populations, considering only ’mean’ matrices (i.e., one matrix per population) with annual time steps. Before the analysis we standardized all MPMs to have λ_0_ = 1 by dividing each matrix element by λ_1_ calculated from the original matrix (see Supporting information S1 for complete description of selection criteria). One hundred fifty-four MPMs were selected, describing two amphibian, 35 bird, 22 bony fish, three insect, 61 mammal, and 31 reptile populations, belonging to 107 species. Generation time was calculated as the mean age of parents at the stable (st)age distribution (Bienvenu & Legendre, 2015) and ranged from 1.1 years to 265.6 years.

### Nonlinear relationships

We added environmental effects to the survival and fertility coefficients. Since some models were stage-structured, we first separated out the two matrices containing these coefficients: Each stage structured projection matrix can be written as **A** = **GS** + **QB** (Vindenes *et al*., 2021). Here **G** and **Q** are matrices describing the stage transition rates of individuals and new offspring, respectively, assumed constant in our analysis. The matrix **B** contains the stage-specific fertility coefficients *f*_*j*_(*z*) on the diagonal and zeroes elsewhere, while the matrix **S** contains stage-specific survival rates *s*_*j*_(*z*) on the diagonal and zeroes elsewhere. For each MPM, we chose a link function for the survival rates *s*_*j*_(*z*) (logistic or loglog link) and a link function for the fertility coefficients *f*_*j*_(*z*) (logistic, loglog, or log link), defining the relationship of **A**(*z*) to the environmental driver *z*. For each scenario we defined different link functions (Fig. 1B-C), where *s*_*j*_(0) and *f*_*j*_(0) corresponded to the values from the standardized MPM in COMADRE. For instance, with a loglog link function, the survival rates are defined as 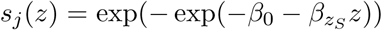, and the parameter *β*_0_ is defined as *β*_0_ = *−* ln(*−* ln(*s*_*j*_(0))). The parameter 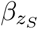 defines the strength of the environmental effect on *s*_*j*_(*z*), and affects the curvature and variance of survival probability in stage *j* (Fig. 1A; Fig. S6 shows survival and fertility coefficients for different 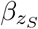 and 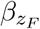 values). Fertility coefficients are defined in a similar way, but here we also defined a maximum MaxF = *M ∗ f*_*j*_(0) with *M* = 2.5 (results for different values of *M* are shown in Supporting information S5), so that the fertility in the mean environment was set as a proportion of the maximum fertility. The values in the mean environment *s*_*j*_(0) and *f*_*j*_(0), defined by the standardized MPM, affect the second derivatives of the link functions (Fig.1A and Fig.S2). A complete description including equations for all link functions and their derivatives is provided in Supporting Information S2 and S7.

To limit the number of scenarios we made the simplifying assumption that survival rates of different (st)ages all have the same value of 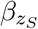, and similarly all fertility coefficients have the same 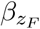. This means that there is always positive covariance among survival rates of different (st)ages and among fertilities of different (st)ages, while covariance between survival and fertility is controlled in our scenarios by the sign of 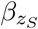 and 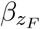. These assumptions are biologically relevant for populations where individuals of different (st)ages live in the same environment, and where survival of different stages and reproduction of different stages are affected similarly by a key driver. A range of other scenarios are also possible but not considered here, such as no covariance among demographic parameters.

### Scenarios

The decomposition of the stochastic growth rate was done under 13 scenarios (Fig. 1C) varying 1) the type of link function defining *s*_*j*_(*z*) and *f*_*j*_(*z*), 2) the covariance between survival and fertility; negative or positive (scenarios 1-8), and by applying 3) special cases of forced buffering, turning off the effect of *z* for certain demographic parameters (thus nonlinearity, variance and covariance of demographic parameters were affected; scenarios 9-13). In the first eight scenarios, effects of *z* were added to survival and fertility of all stages as described above. Four combinations of link functions were tested, including logistic functions for all parameters, loglog link functions for all parameters, and two combinations of log-link function for *f*_*j*_(*z*) with logistic or loglog link functions for *s*_*j*_(*z*). Each of these four combinations was tested using positive or negative covariance between survival and fertility (Fig. 1). In the scenarios of demographic lability with forced buffering, mature stages were defined as all stages equal to or larger than the stage with first non-zero fertility, and immature stages as all stages preceding this stage. Either *s*_*mature*_(*z*) or all *s*_*j*_(*z*) (scenarios 9-11), or all *f*_*j*_(*z*) and *s*_*immature*_(*z*) (scenarios 12-13) were held constant and equal to their value in the mean environment as reported in the standardized COMADRE MPM. We used logistic functions for the other demographic parameters (Fig. 1C). These scenarios reflect different assumptions of demographic lability and buffering within the least or the most influential demographic parameters on population growth, assessed qualitatively depending on the position of the populations along the slow-fast continuum of life histories (Stearns, 1989; Sæther & Bakke, 2000; Gaillard & Yoccoz, 2003). Survival of immature stages and fertility coefficients are assumed to show a higher contribution to population growth in fast-living species, while survival rates of the mature stages are assumed to be more influential for slow life histories.

### Decomposition

For each population in each scenario, we calculated and decomposed the stochastic growth rate ln(λ_*s*_) following equation 3. Since all the MPMs from COMADRE were standardized so that ln(λ_0_) = 0, the stochastic growth rate is a sum of the nonlinearity and the variance-covariance component. The sign of the stochastic growth rate directly reflects whether the fitness effects of environmental variance 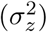 are positive or negative in that population and scenario. All calculations shown in the main text use the value 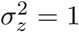, and altering this value only affects the magnitude of the effects (Supporting information S4). In our analyses, 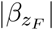 and 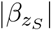 were both set to 0.4 (Fig. 1A; results for other values shown in Supporting information S4).

## Results

### Combined effects of nonlinearity and variance-covariance among demographic parameters

In all scenarios, life histories with short to intermediate generation times (< 10 years) showed consistently stronger fitness responses to environmental variability than slow life histories (Fig. 2-3). Whether these responses are positive or negative, strongly depends on the combined impacts of covariance structure between the (st)age-specific survival rates and fertility coefficients and their curvatures.

**Figure 2.**
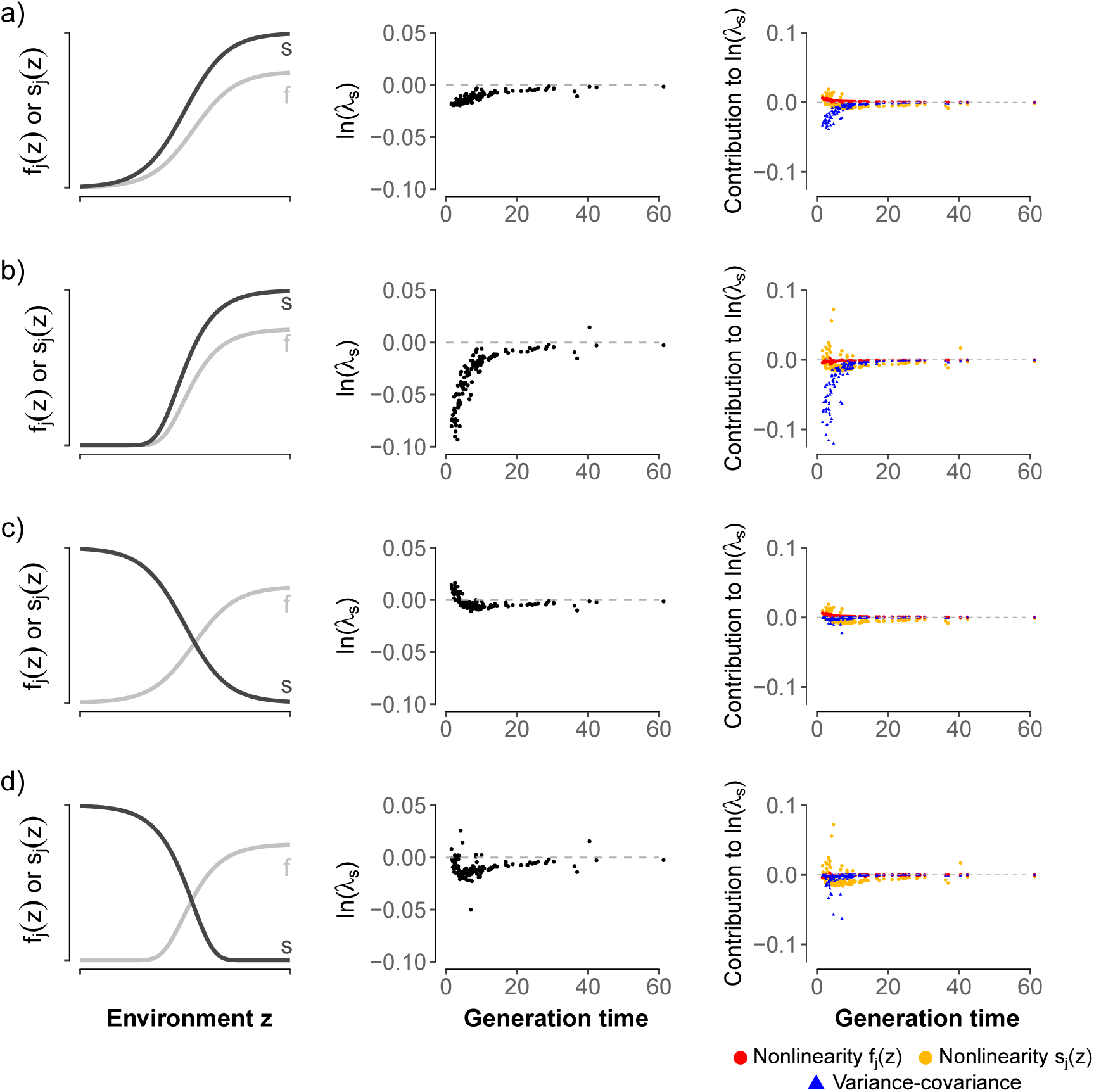
Mid panels: Stochastic growth rate (fitness) ln(λ_*s*_) across generation time, under four scenarios of covariance and link-functions of the demographic parameters. Left panels: Illustration of scenarios, with grey and black lines corresponding to the (st)age-specific survival rates *s*_*j*_(*z*) and fertility coefficients *f*_*j*_(*z*), respectively (functions varied for each stage depending on *s*_*j*_(0) and *f*_*j*_(0); only one function is shown for survival and fertility here). We assumed positive covariance between survival rates of different (st)ages and between the fertilities of different (st)ages. For each scenario and for each population, positive (panels a,b) or negative (panels c,d) covariance between *f*_*j*_(*z*) and *s*_*j*_(*z*) were considered, treating *f*_*j*_(*z*) and *s*_*j*_(*z*) as logistic functions (panels a,c) or loglog link functions (panels b,d) of the environment z. Right panels: Decomposition of ln(λ_*s*_) into main components capturing variance-covariance effects (blue triangles) and lability effects generated by nonlinear responses of *f*_*j*_(*z*) (red circles) and *s*_*j*_(*z*) (orange circles). Results for bony fish populations and populations with generation time > 62 years are not shown (*N*_*MP Ms*_ = 129; see Fig. S14 for all MPMs).

**Figure 3.**
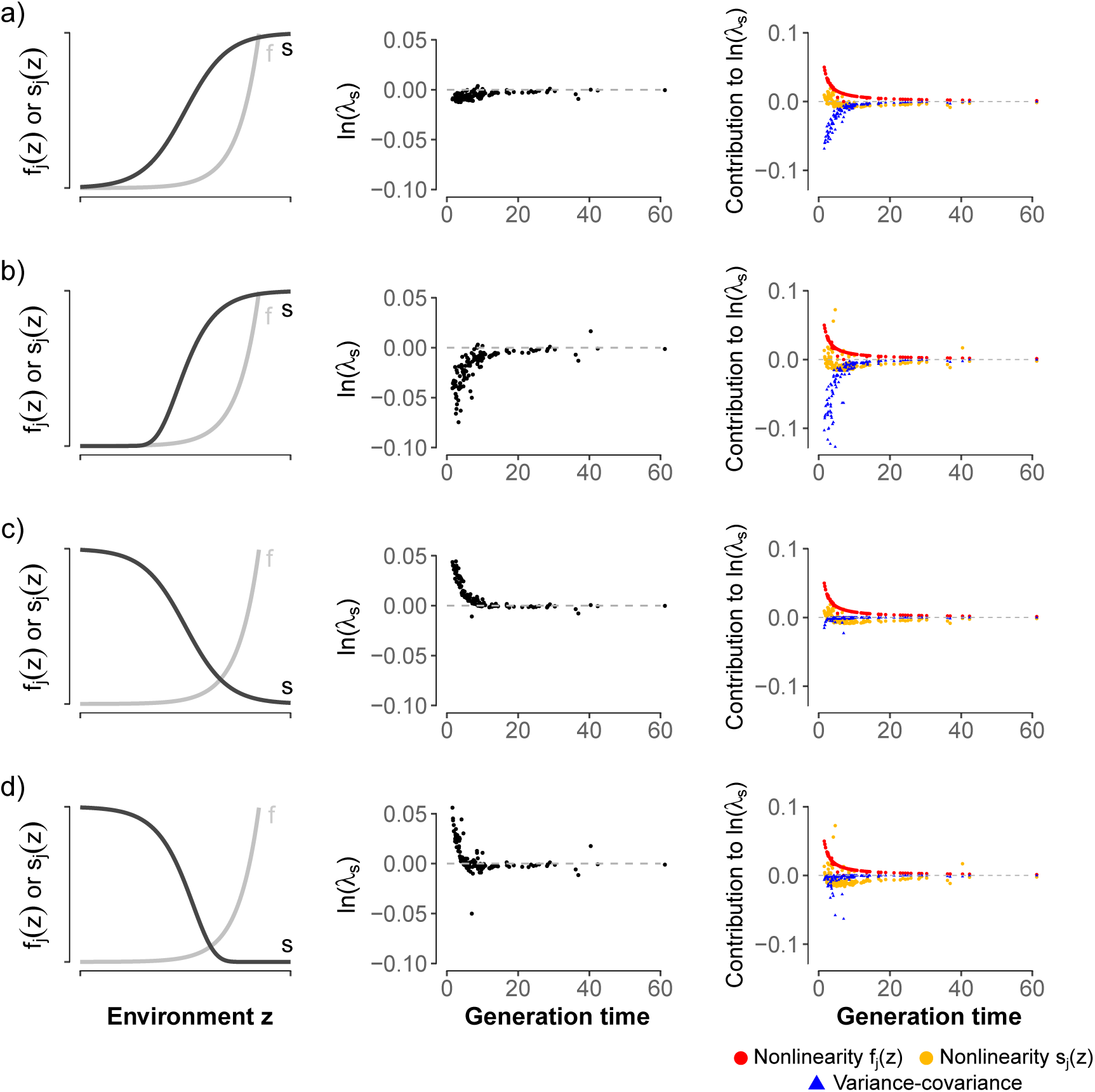
Mid panels: Stochastic growth rate (fitness) ln(λ_*s*_) across generation time, considering positive (panels a,b) or negative (panels c,d) covariance between (st)age-specific survival rates *s*_*j*_(*z*) and fertilities *f*_*j*_(*z*), treating *s*_*j*_(*z*) as logistic (panels a,c) or loglog (panels b,d) link functions of the environment z and *f*_*j*_(*z*) as log link functions. We assumed positive covariance between survival rates of different (st)ages and between the fertilities of different (st)ages. See Fig. 2 for explanation of left and right panels. Results for bony fish populations and populations with generation time > 62 years are not presented (*N*_*MP Ms*_ = 129; see Fig. S15 for all MPMs).

Positive effects of lability on the mean fitness ln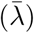 were found mainly among the fast-living species, and positive effects occurred through both survival and fertility (Fig. 2-3). The nonlinearity index D correlated almost perfectly with this nonlinearity component (Spearman coefficient > 0.999 and 0.928 in all scenarios without and with bony fish MPMs, respectively), suggesting that this is a reliable indicator of adaptive lability. However, as fitness ln(λ_*s*_) also depends on the variance-covariance structure of the demographic parameters, this must also be taken into account.

With positive covariance between survival and reproduction, ln(λ_*s*_) was consistently reduced compared to the mean environment, regardless of the type of link functions used (e.g., Fig. 2a, b). In these scenarios, positive nonlinearity components still occurred, but were not sufficient to overcome the negative variance-covariance component. In contrast, populations of fast-living species could show an overall positive fitness ln(λ_*s*_) if survival and fertility covaried negatively (Fig. 2 and 3), although less frequent when *s*_*j*_(*z*) and *f*_*j*_(*z*) were defined by loglog link functions (Fig. 2d). Positive effects were stronger when we used a log-link function for the fertility coefficients, so that they increased exponentially with the environmental driver z leading to strong convexity (Fig. 3c-d). For bony fish MPMs, the signs of the nonlinearity and variance-covariance components were the same as for the other MPMs, but the magnitude was stronger. Here the underlying models from COMADRE showed very high fertility coefficients and low survival rates, yielding extremely high variance in demographic parameters. Under scenarios using loglog link functions for *s*_*j*_(*z*) and/or *f*_*j*_(*z*), the small noise assumption behind our decomposition of ln(λ_*s*_) was violated to a degree where the approximation broke down for these MPMs (Supporting information S6).

### Demographic lability with forced buffering

In these scenarios, some survival probabilities and fertility coefficients were kept constant and buffered, while others were allowed to vary. The identity of labile demographic parameters, together with the position of the species along the slow-fast continuum, affected each fitness component and their combined impact on fitness (Fig. 4). When lability in all survival rates *s*_*j*_(*z*) or in only the mature stages *s*_*mature*_(*z*) was combined with a constant fertility (Fig. 4a-b), only the fastest-living species showed a positive ln(λ_*s*_). This positive fitness resulted from a positive nonlinearity effect of survival rates and a low negative variance-covariance effect, reflecting buffering. When lability in fertility *f*_*j*_(*z*) and survival rates of the immature stages *s*_*immature*_(*z*) was combined with constant survival rates in all stages or mature stages, positive values of ln(λ_*s*_) were also detected, especially when immature survival rates and reproduction covaried negatively (Fig. 4c-e).

**Figure 4.**
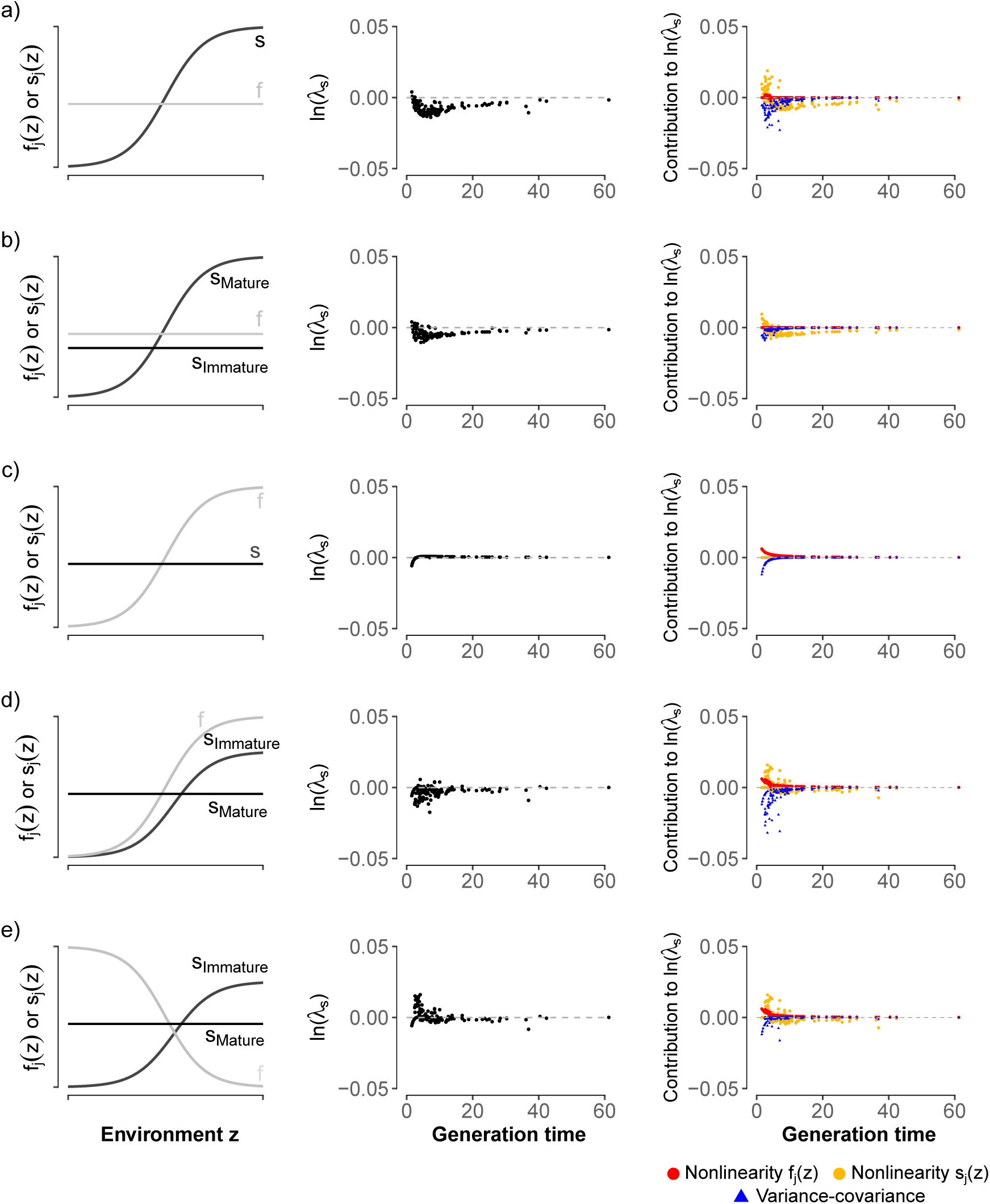
Results from scenarios of forced buffering assuming demographic lability only in (a) (st)age-specific survival rates, (b) survival rates of the reproductive stages only, (c) (st)agespecific fertilities and (d-e) fertilities and survival rates of the immature stages. For each scenario, the long term fitness ln(λ_*s*_) and its main components reflecting variance-covariance effects (blue triangles) and lability effects due to nonlinearity of *f*_*j*_(*z*) (red circles) and *s*_*j*_(*z*) (orange circles) are plotted against generation time (mid and right panels; see Fig. S16 for all MPMs). See Fig. 2 for explanation of left panel.

In contrast, for intermediate and slow-living species, labile survival rates of the reproductive stages *s*_*mature*_(*z*) combined with constant fertility *f*_*j*_(*z*) and constant survival of immature stages *s*_*immature*_(*z*) (Fig. 4b) always produced negative nonlinearity components, and very small negative variance-covariance components, leading to an overall negative ln(λ_*s*_). The scenarios of lability in fertility coefficients combined with constant (st)age-specific survival rates or in only the mature stages (Fig. 4c-e) showed a weak negative variance-covariance component while the nonlinearity component was zero or slightly positive, leading to overall fitness ln(λ_*s*_) having values close to zero. In other words, constant (st)age-specific survival rates associated with labile fertility coefficients have a stabilizing effect on ln(λ_*s*_) of slow life histories (generation time > 10 years; Fig. 4c VS Fig. 4a-b).

## Discussion

This study emphasizes the importance of considering explicit links between environmental drivers and demographic parameters to understand the effects of environmental variability on fitness, as these links allow effects on nonlinearity to be quantified. We extended Tuljapurkar’s approximation of the stochastic growth rate to incorporate effects of nonlinearity in demographic parameters. We also defined a nonlinearity index to measure the overall nonlinearity in a given life history, reflecting the potential for positive fitness effects of environmental variability. Our decomposition of the stochastic growth rate into nonlinearity and variance-covariance components creates a new framework to study their joint impacts on fitness, expanding earlier theory focusing mainly on buffering through the variance-covariance component. Applying this decomposition across a range of scenarios and life histories, we identified the faster-living species as the most responsive to environmental fluctuations, both through the nonlinearity and variance-covariance components. Positive fitness values were only found when positive nonlinearity components were combined with negative covariance between survival and fertility, leading to a smaller negative variance-covariance component. In scenarios with some demographic parameters being constant (forced buffering), lability in both the least and the most expected influential demographic parameters were found to benefit fitness to some extent, but mainly for short-lived species. Our decomposition provides a step forward in our understanding of potential adaptations to environmental variability in a wide range of life histories, and stresses the importance of characterising both nonlinearity and covariance structure of demographic parameters with respect to key environmental drivers. Our framework is also useful for predicting population responses to increased variability under global change.

### Lability and buffering in fast vs. slow life histories

Several studies have shown evidence that populations located at the fast end of the slow-fast continuum are more sensitive to changes in the different components of climate change. These populations tend to respond more strongly to changes in climate drivers (e.g., Compagnoni *et al*. 2021), to environmental variability (e.g., Dalgleish *et al*. 2010; Drake 2005; Koons *et al*. 2009; Morris *et al*. 2008, but see Le Coeur *et al*. 2021; Santos *et al*. 2021), to shifts in temporal autocorrelation in the environment (e.g., Paniw *et al*. 2018), and to shifts in the correlation structure of demographic parameters (Iles *et al*., 2019). In line with these previous studies, we found that populations of faster-living species have larger absolute values of both nonlinearity and variance-covariance components of fitness in a stochastic environment compared to those of slow living-species. On one hand, fast-living species are more vulnerable to environmental fluctuations due to higher negative variance-covariance components, as reported in previous studies (e.g., Dalgleish *et al*. 2010; Morris *et al*. 2008). On the other hand, they have the largest potential for adaptive lability through convex demographic responses. Our results show that a positive nonlinearity component can overcome the negative variance-covariance and lead to increased fitness especially when there is a negative correlation between fertility and survival. We found that the nonlinearity index D is a reliable predictor of the nonlinearity component of the stochastic growth rate (eq. 3).

A majority of studies have focused on effects of the variance-covariance component alone, without explicit reference to the underlying environmental drivers, even though other studies (Drake, 2005; Henden *et al*., 2008; Koons *et al*., 2009) highlighted the potentially critical importance of including such links. Our results support this conclusion, and show that the total impact of environmental fluctuations on the fitness of structured populations may be either positive or negative if nonlinear demographic responses are present (eq. 3). With explicit links, where some are convex, positive fitness responses are possible, but we highlight that the net effect also depends on the variance-covariance component and the type of link functions. Evidence of convex relationships between demographic parameters or underlying vital rates and key environmental drivers is still limited for natural populations, due to data limitation or *a priori* linear assumptions in the statistical models. Our study highlights the need for empirical research to determine more systematically the shape and curvature of demographic parameter responses to accurately predict fitness responses to environmental variance. Quantifying the relationships between environmental drivers and all demographic parameters remains, however, a statistical challenge for wild populations (e.g., separating link functions; Gill, 2001) and requires long-term monitoring data (see Lee, 2017 for an alternative method to study nonlinearity in the growth rate response to an environmental driver with discrete levels). This highlights the need to continue and increase the ongoing collection of demographic data.

The decomposition of the stochastic growth rate considers nonlinearity and variance-covariance of demographic parameters, which in turn are functions of underlying vital rates. For instance, fertility depends on both fecundity and survival of offspring or parents, depending on the census of the matrix model. Studies applying the method for specific empirical systems should carefully consider how the demographic parameters depend on lower-level parameters as functions of environmental drivers. Our qualitative conclusions on demographic buffering and lability across generation time are general, but quantitative differences are likely present for instance for models based on prevs. post-reproductive census, when environmental effects arise through lower-level parameters. This presents an interesting area for future research using the decomposition.

### Role of temporal covariance between (st)age-specific demographic parameters

While negative covariance between demographic parameters could arise from life history tradeoffs (Stearns, 1989) or opposite responses to the same environmental driver, positive covariances between these parameters are just as likely to occur in a population. Previous theoretical work has shown that positive covariance enhances the variance in population growth while negative covariance reduces it (Tuljapurkar, 1982, 1990). Our results are in line with this result, showing reduced negative variance-covariance component when survival and fertility covaried negatively compared to positively.

Interestingly, there is no general consensus on the degree to which positive or negative covariance in demographic parameters are more common in the wild, nor if the sign, magnitude or type of (st)age-specific demographic parameters involved correlate with the position of a species along the fast–slow continuum (but see a recent comparative study, Fay *et al*., 2022). From empirical studies, positive covariances have been reported predominantly in long-lived species (e.g., Dahlgren *et al*., 2016; Rotella *et al*., 2012; van de Pol *et al*., 2010) with substantial (e.g., Coulson *et al*., 2005) or weak (e.g., Altwegg *et al*., 2007; Compagnoni *et al*., 2016; Johnson *et al*., 2010) effects on fitness. In contrast, negative covariances were less often detected (Fay *et al*., 2022), with often small effects on ln(λ_*s*_). To our knowledge, relatively few studies have specifically addressed this question among species towards the fast-end of the continuum.

In our scenarios, we assumed a perfect, positive temporal covariance between (st)age-specific survival rates and between (st)age-specific fertilities, respectively, but positive or negative covariances between survival and fertility. While these assumptions on the direction of covariance between stages and type of demographic parameters are plausible, they are strong in terms of magnitude, and a main environmental driver is unlikely to explain all of the (co)variance in demographic parameters. Our results may therefore overestimate the magnitude of the variance-covariance component in the decomposition, compared to wild populations where correlations are likely not perfect. Even though we assumed perfect correlation, we found that variance-covariance had negligible effects on fitness of slow-living populations, reflecting a large degree of buffering in these species. For fast-living species, covariance had contrasting effects on the fitness components. These effects were strengthened in scenarios where link functions implied more asymmetric relationships between demographic parameters and environmental driver.

### Demographic lability and buffering of different demographic parameters

The set of scenarios combining lability in some demographic parameters with forced buffering in others, yielded insights into possible demographic strategies along the slow-fast continuum. While different predictions have been made as to which demographic parameters should be selected for lability (Hilde *et al*., 2020; McDonald *et al*., 2017), we found that demographic lability in the demographic parameters assumed to be the least (*f*_*j*_(*z*) and/or *s*_*immature*_(*z*)) or most (*s*_*mature*_(*z*)) important to fitness, could both lead to enhanced fitness in many fast-living life histories due to positive nonlinearity components and reduced variance-covariance components. However, such positive effects on fitness were stronger and more prevalent with lability in both fertility and survival of the immature stages (the most influential in fast life histories). In contrast, for slow-living life histories, lability in the survival rates of mature stages, believed to have the highest impact on fitness led to negative effects on fitness due to negative nonlinearity components. Selection for a reduction in variance in (and in positive covariance between) the demographic parameters that contribute the most to fitness, combined with other parameters varying more freely, as stated by the demographic buffering hypothesis, seems likely for slow-living species, at least those with a similar animal life history as in our analysis.

Labile and buffered demographic parameters in our scenarios were qualitatively assigned based on expectations from the demographic buffering and life history theories (Stearns, 1989; Sæther & Bakke, 2000; Gaillard & Yoccoz, 2003). This simple categorization, while accurate for some life histories, may be different for other populations with the same generation time. Further insights would require differentiating labile and buffered (st)age-specific demographic parameters and underlying vital rates in a population based on elasticities of the growth rate in the mean environment.

In conclusion, this study provides a comprehensive framework for assessing the contributions of demographic lability and buffering on fitness of any given population. Positive effects of environmental fluctuations on fitness are only possible to detect if we account for the impacts of nonlinear relationships between demographic parameters and environmental drivers. Our decomposition of the stochastic growth rate into components of nonlinearity and variance-covariance provides a tool to quantify their relative impacts in different life histories and scenarios, and is easily applicable for other study systems and scenarios not considered here. Across the slow-fast continuum of animal life histories, faster-living species have the largest potential for using demographic lability as an adaptive response to variability, while demographic buffering is a main adaptive response in slow-living species. These findings have important implications for predicting population and species responses to changes in environmental fluctuations under climate change and other anthropogenic impacts.

## Text box 1: Glossary

### Stochastic population growth rate - ln(λ_s_)

The long-term rate of population growth on a logarithmic scale, a measure of fitness in a stochastic density-independent environment.

### Growth rate in the mean environment - ln(λ_0_)

dominant eigenvalue of the projection matrix in the mean environment (z=0) **A**(**0**) on a logarithmic scale.

### Mean growth rate – ln 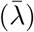

dominant eigenvalue of the mean projection matrix across variable environments 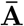 on a logarithmic scale.

### Demographic lability / labile demographic parameter

A labile demographic parameter fluctuates with temporal variation in environmental conditions. The relationship between a labile demographic parameter and the environment (e.g., a key environmental driver) can be convex, concave or linear, so that the average value of this demographic parameter in a variable environment becomes >, <, or = to the demographic parameter estimated in the mean environment (z=0), respectively. The same definition applies to labile vital rates (e.g., survival, fecundity, transition).

### Adaptive demographic lability (demographic lability hypothesis)

selection for demographic parameters to track environmental fluctuations that leads to an overall increased fitness, ln(λ_*s*_). Increase in ln(λ_*s*_) occurs when an increase in the demographic parameter means due to convexity in their responses leads to a shift in the arithmetic mean of annual population growth rates ln 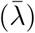, that overcomes the negative effect of temporal variance in the annual population growth rates (variance-covariance component *σ*^2^). This hypothesis relies on the assumption that the nonlinearity index *D* (defined below) is positive.

### Nonlinearity index (D)

*D* measures the total effect of nonlinearity of demographic parameters in a life history, and is a key component to describe the nonlinearity component of the fitness decomposition (equation 3). This index corresponds to the sum over all (st)ages of the second derivatives of the demographic parameters (depending on vital rates) in the mean environment (z=0), weighted by the sensitivities of λ_0_ to the corresponding demographic parameters (matrix elements). When positive (/negative), *D* is an indicator of adaptive (/non-adaptive) lability through overall positive (/negative) contributions from convexity (/concavity) of the demographic parameters. Adaptive lability can create a positive overall effect of environmental variability if *D* is positive and the negative effects of increased variance-covariance of the demographic parameters are not too large (see equation 3).

### Demographic buffering / buffered demographic parameters

Low variance of a demographic parameter in response to temporal variation in the environmental variable *z*. A more flat relationship between the demographic parameter and the environment *z* leads to such low parameter variance, and to the mean demographic parameter in the variable environment remaining approximately equal to demographic parameter value in the mean environment (*z*=0). The same definition applies to buffered vital rates (e.g., survival, fecundity, transition).

### Adaptive demographic buffering (demographic buffering hypothesis)

The prediction that natural selection should favour a reduction in variance of the demographic parameters with the strongest influence on fitness in the mean environment, reducing the variance-covariance component *σ*^2^ and leading to an overall stable or increased fitness in variable environments. The assumption that ln 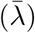 is not affected by environmental variance 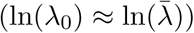, is often made for this hypothesis.

