## Supporting information for "Life history adaptations to fluctuating environments: Combined effects of demographic buffering and lability of demographic parameters"

**S1 - Matrix population models and selection criteria** **2**

**S2 - Nonlinear functions relating demographic parameters to an explicit environment driver** **5**

**S3 - Extending Tuljapurkar's approximation of the stochastic growth rate to include nonlinear effects** **9**

**S4 - Simulations to test accuracy of the decomposition - comparison of different environmental variance levels $\sigma_z^2$ and environmental strength $\beta_z$** **11**

**S5 - Comparison of threshold values in (st)age-specific fertility coefficients** **20**

**S6 - Results including matrix population models of bony fishes** **23**

**S7 - R code: decomposition of the stochastic population growth rate** **29**

### S1 - Matrix population models and selection criteria

To explore life history variation in the stochastic growth rate and its main components, we used age- and stage-structured matrix population models (MPMs) from the COMADRE Animal Matrix Database (v.4.20.5; Salguero-Gómez *et al.* (2016)). This database contains over 3360 matrix population models for more than 400 species. We selected matrix population models given some selection criteria and let these projection matrices represent the matrices in the mean environment. Only mean matrices (i.e, one matrix per population) with annual time steps of unmanipulated and free-ranging populations were considered.

We used the following R command for selection:

```
subset(comadre, Composition == "Mean - a mean matrix was calculated by
                 averaging across more than one population studied over a single time
                     period, for example, over one year"
               & MatrixFec =="1"
               & StudiedSex!= "M/F - Males and females separately in the same
                   population matrix model/F"
               & MatrixTreatment == "Unmanipulated"
               & ProjectionInterval == 1
               & MatrixCaptivity == "W - Wild: study in natural conditions"
               & SurvivalIssue <1)
```

As a second step, we standardized all matrices to $\lambda_0 = 1$ by dividing each matrix element by $\lambda_1$, the deterministic growth rate calculated from the original matrix ($\lambda_1 \in$ [0.42-4.64] with a mean of 1.09). In some cases of very low original $\lambda_1$-values this resulted in survival and transition probabilities exceeding 1, and these models were discarded from the analysis. This standardization allows reliable comparisons between populations, avoiding confounding effects associated with declining or increasing populations on $ln(\lambda_s)$ and population extinction during stochastic simulations.

Overall, a total of 154 MPMs describing two amphibian, 35 bird, 22 bony fish, three insect, 61 mammal and 31 reptile populations, belonging to 107 species were included in the analysis.

List of "Matrix ID" used in the simulation framework:
240307, 240308, 240309, 240310, 240646, 240647, 240648, 240651, 249118, 249119, 249122, 249133, 249159, 249160, 249182, 249191, 249214, 249248, 249249, 249250, 249251, 249272,

249274, 249286, 249292, 249332, 249376, 249410, 249411, 249412, 249464, 249504, 249506, 249512, 249524, 249525, 249605, 249619, 249620, 249621, 249625, 249674, 249734, 249757, 249758, 249759, 249810, 249811, 249812, 249837, 249851, 249871, 249876, 249877, 249880, 249881, 249882, 249897, 249909, 249921, 249922, 248391, 248392, 248398, 248404, 248405, 248406, 248413, 248433, 248441, 248446, 248451, 248452, 248454, 248473, 248478, 248515, 248518, 248552, 248553, 248575, 248615, 248616, 248631, 248635, 248688, 248696, 248700, 248702, 248715, 248718, 248722, 248763, 248770, 248787, 248818, 249020, 249049, 249052, 250003, 250023, 250027, 250029, 250037, 250041, 250047, 250057, 250058, 250060, 250088, 250094, 250114, 250115, 250116, 250118, 250120, 250122, 250123, 250127, 250128, 250129, 250130, 250131, 250132, 250133, 250134, 250158, 250163, 250164, 250168, 248236, 248240, 248046, 248048, 248057, 248091, 248104, 248109, 248111, 248128, 248150, 248151, 248152, 248154, 248155, 248156, 248166, 248169, 248173, 248178, 248186, 248220, 248221, 248222.

### Selected MPMs along the slow-fast continuum

We checked that the selected MPMs ($N_{MPMs} = 154$) were representative of a wide range of life histories we see in nature. To that end, generation time and the degree of iteroparity (based on the Demetrius' entropy) were estimated for these matrices and compared to a wider range of Comadre matrices (Selection 1: $N_{MPMs} = 557$; selection 2: $N_{MPMs} = 692$). Demetrius' entropy was estimated using the Rage package (Jones *et al.*, 2021) and generation time was calculated as the mean age of parents at the stable (st)age distribution (Caswell, 2001). The Demetrius' entropy could not be estimated for 45 out of the 154 MPMs. We found that our MPM selection was a representative sample of the life histories gathered in the large COMADRE Animal Matrix Database (Fig. S1).

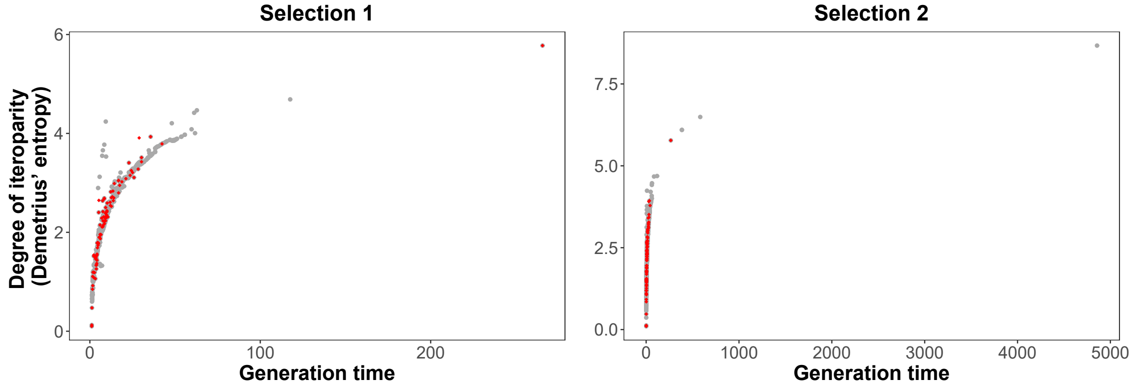


Figure S1: Relationship between Demetrius' entropy and generation time estimated from our selected MPMs (red triangles) and compared to two wider selection of MPMs. The selected MPMs are representative of the life histories gathered in COMADRE Animal Matrix Database.

We used the following R command for selection 1:

```
subset(comadre, MatrixFec =="1"
              & StudiedSex!= "M/F - Males and females separately in the same
                  population matrix model/F" & MatrixTreatment == "Unmanipulated"
              & ProjectionInterval == 1
              & MatrixCaptivity == "W - Wild: study in natural conditions"
              & SurvivalIssue <1)
```

And for selection 2:

```
subset(comadre, MatrixFec =="1"
              & StudiedSex!= "M/F - Males and females separately in the same
                  population matrix model/F" & ProjectionInterval == 1
              & SurvivalIssue <1)
```

### S2 - Nonlinear functions relating demographic parameters to an explicit environment driver

For each matrix population model and for each scenario, we chose a link function for the survival rates $s_j(z)$ (logistic or loglog link) and a link function for the fertility coefficients $f_j(z)$ (logistic, loglog, or log link) to determine the type of relationship to the main environmental driver $z$. We assume that the environmental driver $z$ is a stochastic variable (independent and identically distributed) with mean E[$z$]=0 and variance $\sigma_z^2$= 0.1, 0.5, 1, 1.5 or 2. For the simulation approach, z $\sim \mathcal{N}(0, \sigma_z^2)$.

We chose these functions (logistic, loglog or log link functions) because they represent a broad range of biologically plausible relationships with a key environmental driver (resource, predation, weather). If the slope term $\beta_z$ is relatively high, it induces strong effects on the variance and curvatures of the demographic parameters using the loglog link functions and strong convex and exponential relationships using the log link function for the fertility coefficients. We are confident that the three types of relationships considered covered small to moderate magnitude of lability and variance in the demographic parameters. Note that altering the $\beta_z$ values is equivalent to modifying the variance $\sigma_z^2$ in terms of increasing the effects of environmental fluctuations, but by including $\sigma_z^2$ the link to the environmental driver $z$ is made explicit. Functions are defined below and first and second derivatives of each function are given in Supporting information 7 (R code).

- **Stage-specific fertility coefficients as logistic functions of *z*,** $f_j(z)$:

  Fertility coefficient of age class or stage $j$ is defined as a logistic function of $z$:

$$f_j(z) = \frac{MaxF}{1 + \exp(-\beta_0 - \beta_{z_F} * z)} \tag{1}$$

  where MaxF (maximum fertility)= M * $f_j(0)$ with M= 1.5, 2.5 or 3. $f_j(0)$ denotes the fertility coefficient of (st)age $j$ in the COMADRE matrix (z=0), limiting the range of values to $f_j(z) \in$ [0, M * $f_j(0)$]. $\beta_0$ = logit(1/M) and $\beta_{z_F}$ determines the steepness of the slope, i.e., the strength of the environmental effect on $f_j(z)$.

- **Stage-specific fertility coefficients as loglog link functions of *z*,** $f_j(z)$**:**

$$f_j(z) = MaxF * \exp(-\exp(-\beta_0 - \beta_{z_F} * z) \tag{2}$$

where MaxF = M * $f_j(0)$ with M= 1.5, 2.5 or 3. $f_j(0)$ denotes the fertility coefficient of (st)age $j$ in the COMADRE matrix, $\beta_0$ = -ln(-ln(1/M)) and $\beta_{z_F}$ determines the steepness of the slope.

- **Stage-specific fertility coefficients as log link functions of $z$, $f_j(z)$:**

$$f_j(z) = \exp(\beta_0 + \beta_{z_F} * z) \text{ where } \beta_0 = \ln(f_j(0)). \tag{3}$$

Survival rates for each age class or stage $j$ were defined as a logistic or a loglog link function of $z$ (Fig. 1 in the main body of the manuscript and Fig. S2 below).

- **Stage-specific survival rates $s_j$ as logistic functions of $z$:**

$$s_j(z) = \frac{1}{1 + \exp^( -\beta_0 - \beta_{z_S} * z)} \text{ where } \beta_0 = \text{logit}(s_j(0)) \tag{4}$$

$\beta_{z_S}$ corresponds to the strength of the environmental effect on $s_j(z)$.

- **Stage-specific survival rates $s_j$ as loglog link functions of $z$:**

$$s_j(z) = \exp(-\exp(-\beta_0 - \beta_{z_S} * z) \text{ where } \beta_0 = \text{-ln(-ln}(s_j(0)) \tag{5}$$

In our analysis, we assumed that survival rates of different (st)ages have the same value of $\beta_{z_S}$, and similarly all fertility coefficients have the same $\beta_{z_F}$. This means that the values for the mean environment $s_j(0)$ and $f_j(0)$, defined by the COMADRE matrix, control the curvature of the demographic parameter functions around the mean environment and restrict how large the variance in each demographic parameter can be. Figure S2 illustrates how the curvature change with different $s_j(0)$ and $f_j(0)$ means.

There are a number of alternatives when it comes to simulating variable demographic parameters, with no clear recommendations. Dividing the variance by the maximum variance (or similarly the CV by the maximum CV; Hilde *et al.*, 2020), or using variance-stabilizing transformation such as logit and $arcsin(\sqrt{p})$ following Link & Doherty (2002) have both been used. However, as Link & Doherty (2002) made clear in their paper, the choice should ideally depend on the relationship between the variance and the mean, a relationship which is still not well established empirically. Moreover, the relationships derived in Link & Doherty (2002) are based on using the first-order delta method, and do not work well when the variances are not small, or the proportions close to 0 or 1 (Fig. S3; see also Warton & Hui, 2011 about the arcsine transformation).

The relationships between the variance of rates (proportions) and the coefficients $\beta_z$ used in the simulations are displayed in Fig. S3. It shows that if one had used the standardization by the maximum variance, our parameterization would have worked relatively well, except for values close to 0 and 1 for which there is no clear solutions supported both by empirical observations of the relationship mean-variance and theoretical solutions adjusting for this relationship.

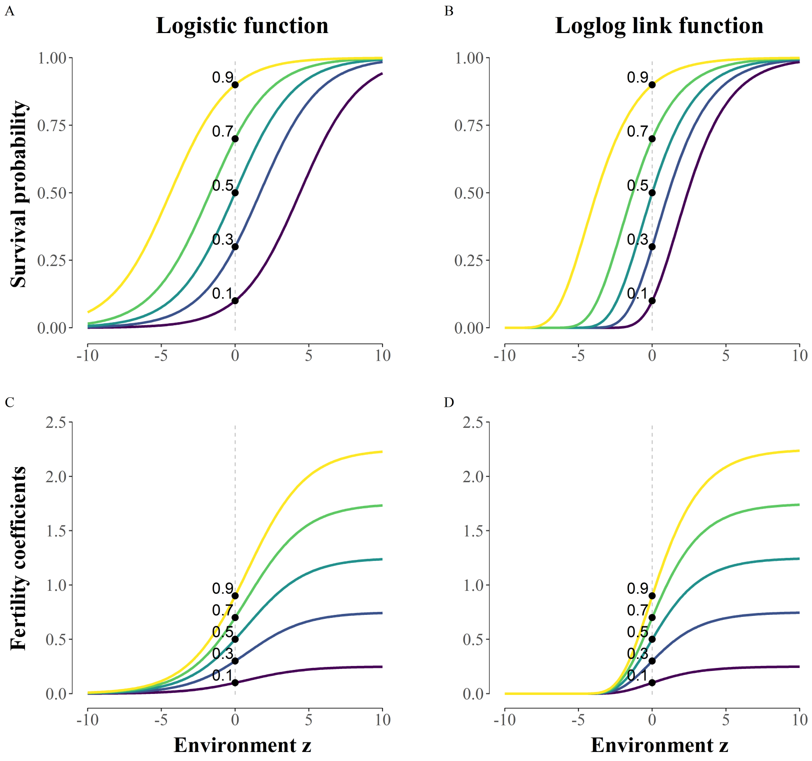


Figure S2: Survival rates (A-B) and fertility coefficients (C-D) as logistic (left plot) and loglog link functions (right plot) of the environment z, with $\beta_{z_S}$=$\beta_{z_F}$=0.5 (high environmental variability scenario) and $MaxF = 2.5 * f_j(0)$. Here, $f_j(0)$ and $s_j(0)$ equal 0.1, 0.3, 0.5, 0.7, 0.9.

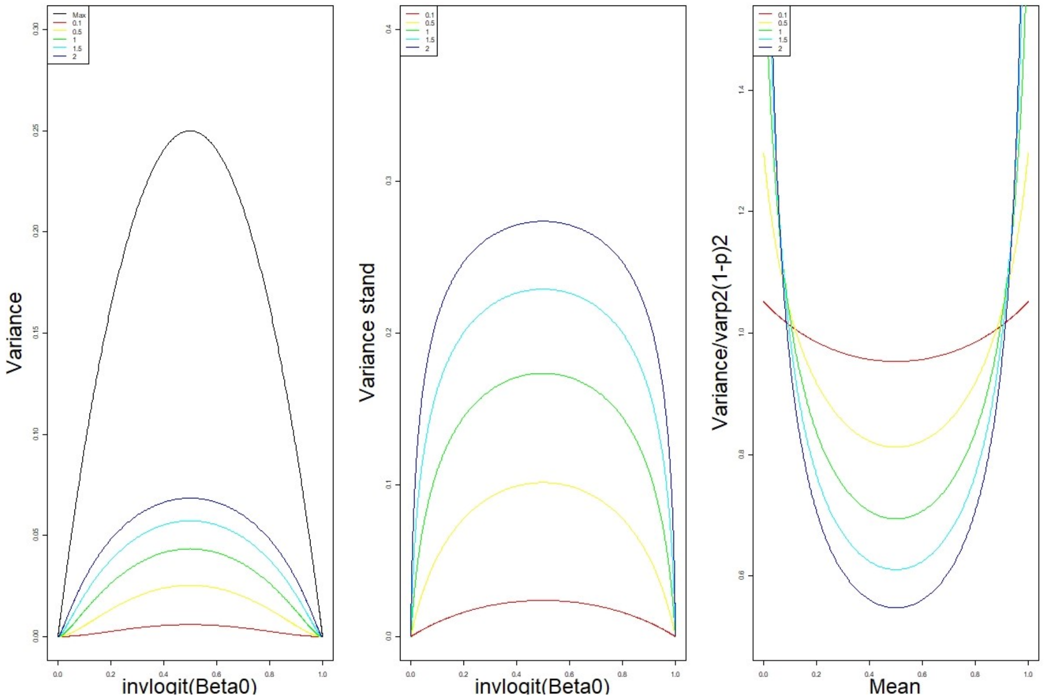


Figure S3: **Left panel:** Relationship between the variance of proportions and the assumed variance ($\beta_z$ in the text) on the logit scale. The max(imum) variance is given by p/(1-p) where p is the average proportion, and $\beta_0$ is the mean on the logit scale. **Middle panel:** Variances standardized by dividing by the maximum variance p(1-p). **Right panel:** Variance divided by the approximated variance calculated using the first-order delta method. The approximation works well for small variance (0.1). The ratio is close to 1 but does not work well for large variances.

### S3 - Extending Tuljapurkar's approximation of the stochastic growth rate to include nonlinear effects

In this appendix we show how to approximate the stochastic growth rate $r_S = \ln \lambda_s$ for a stage structured population where the demographic parameters are possibly nonlinear functions of an environmental variable $z$, with mean $E[z] = 0$ and variance $\mathrm{Var}(z) = \sigma_z^2$. Let $\mathbf{A}(z)$ describe the projection matrix where elements are functions of $z$. Then $\mathbf{A}(0)$ is the matrix in the mean environment, $\mathbf{A}'(0)$ contains the first derivatives of each element of this matrix with respect to $z$, measured in the mean environment, and $\mathbf{A}''(0)$ contains the second derivatives. From the matrix $\mathbf{A}(0)$ we calculate the mean growth rate $\lambda_0$ (dominant eigenvalue), the stable (st)age structure $\mathbf{u}$ (right eigenvector with respect to $\lambda_0$), and the reproductive values $\mathbf{v}$ (left eigenvector associated with $\lambda_0$), scaled so that $\mathbf{vu} = 1$. From these in turn we define the sensitivity matrix $\mathbf{S}$ with elements $S_{ij} = v_i u_j$.

Applying a second order Taylor approximation to each element of $\mathbf{A}(z)$, we can write the mean of the stochastic $k \times k$ projection matrix $\mathbf{A}(Z)$ as $\mathrm{E}[\mathbf{A_t}] \approx \mathbf{A}_0 + \frac{\sigma_z^2}{2}\mathbf{A}''(0)$. Then we can decompose the stochastic matrix for each time step $t$ as $\mathbf{A}_t \approx \mathbf{A}_0 + \frac{1}{2}\mathbf{A}''(0) + \varepsilon_t$, where $\varepsilon_t$ is the matrix containing the stochastic deviations from $E[\mathbf{A}_t]$, with mean 0.

We use this decomposition of the stochastic matrix to write the dynamics of the total reproductive value $V_t = \mathbf{vn}_t$ as

$$V_{t+1} = \lambda_0 V_t + \frac{\sigma_z^2}{2}\mathbf{vA}''(0)\mathbf{n}_t + \mathbf{v}\varepsilon_t\mathbf{n}_t$$

Using the approximation $\mathbf{n}_t \approx \mathbf{u}V_t$ we get

$$\begin{aligned} V_{t+1} &= \lambda_0 V_t + \frac{\sigma_z^2}{2}\mathbf{vA}''(0)\mathbf{u}V_t + \mathbf{v}\varepsilon_t\mathbf{u}V_t \\ &= \lambda_0 \left(1 + \frac{\sigma_z^2}{2\lambda_0}\mathbf{vA}''(0)\mathbf{u} + \frac{1}{\lambda_0}\mathbf{v}\varepsilon_t\mathbf{u}\right) V_t \end{aligned}$$

Note that $\mathbf{vA}''(0)\mathbf{u} = D$ is what we define in the main text as the 'Nonlinearity index'. Taking the logarithm of each side we get

$$\ln V_{t+1} - \ln V_t \approx \ln \lambda_0 + \ln\left(1 + \frac{1}{\lambda_0}\left[\mathbf{v}\varepsilon_t\mathbf{u} + \frac{\sigma_z^2}{2}\mathbf{vA}''(0)\mathbf{u}\right]\right).$$

Using a second order Taylor approximation of the last term, and taking the expectation of this part, we get

$$\begin{aligned}
\mathrm{E}\left[\ln\left(1+\frac{1}{\lambda_0}\left[\mathbf{v}\varepsilon_t\mathbf{u}+\frac{\sigma_z^2}{2}\mathbf{v}\mathbf{A}''(0)\mathbf{u}\right]\right)\right] &\approx \frac{\sigma_z^2}{2\lambda_0}\mathbf{v}\mathbf{A}''(0)\mathbf{u}-\frac{1}{2\lambda_0^2}\mathrm{Var}\left(\mathbf{v}\varepsilon_t\mathbf{u}\right)-\frac{\sigma_z^4}{8\lambda_0^2}\left(\mathbf{v}\mathbf{A}''(0)\mathbf{u}\right)^2\\
&=\frac{\sigma_z^2}{2\lambda_0}\mathbf{v}\mathbf{A}''(0)\mathbf{u}\left[1-\frac{\sigma_z^2}{4\lambda_0}\mathbf{v}\mathbf{A}''(0)\mathbf{u}\right]-\frac{1}{2\lambda_0^2}\mathrm{Var}\left(\mathbf{v}\varepsilon_t\mathbf{u}\right).
\end{aligned}$$

Putting the elements together, the approximation of the stochastic growth rate is

$$\begin{aligned}
r_S &= \mathrm{E}[\ln V_{t+1}-\ln V_t|V_t]\\
&\approx \ln\lambda_0+\frac{\sigma_z^2}{2\lambda_0}\mathbf{v}\mathbf{A}''(0)\mathbf{u}\left[1-\frac{\sigma_z^2}{4\lambda_0}\mathbf{v}\mathbf{A}''(0)\mathbf{u}\right]-\frac{1}{2\lambda_0^2}\mathrm{Var}\left(\mathbf{v}\varepsilon_t\mathbf{u}\right).
\end{aligned}$$

Using a second order Taylor expansion of the variance / covariance matrix (and that $\mathrm{Var}(Z^2) = E[Z^4]-E[Z^2]^2 = 3\mathrm{Var}(Z)^2-\mathrm{Var}(Z)^2 = 2\mathrm{Var}(Z)^2 = 2\sigma_z^4$) we can write the variance term as

$$\begin{aligned}
\mathrm{Var}\left(\mathbf{v}\varepsilon_t\mathbf{u}\right) &= \sum_{ij}\sum_{kl}\frac{d\lambda}{dA_{ij}}\frac{d\lambda}{dA_{kl}}\mathrm{Cov}(A_{ij},A_{kl})\\
&\approx \sigma_z^2\sum_{ij}\sum_{kl}\frac{d\lambda}{dA_{ij}}\frac{d\lambda}{dA_{kl}}A'_{ij}(0)A'_{kl}(0)+\frac{\sigma_z^4}{2}\sum_{ij}\sum_{kl}\frac{d\lambda}{dA_{ij}}\frac{d\lambda}{dA_{kl}}A''_{ij}(0)A''_{kl}(0)\\
&= \sigma_z^2 B+\frac{\sigma_z^4}{2}C.
\end{aligned}$$

Inserting this as well as the nonlinearity index $D$ in the equation for the stochastic growth rate, we get

$$\begin{aligned}
r_S &\approx \ln\lambda_0+\frac{\sigma_z^2}{2\lambda_0}D-\frac{\sigma_z^4}{8\lambda_0^2}D^2-\frac{\sigma_z^2}{2\lambda_0^2}B-\frac{\sigma_z^4}{4\lambda_0^2}C\\
&= \ln\lambda_0+\frac{\sigma_z^2}{2\lambda_0}D\left(1-\frac{\sigma_z^2}{4\lambda_0}D\right)-\frac{\sigma_z^2}{2\lambda_0^2}\left(B+\frac{\sigma_z^2}{2}C\right)\\
&\approx \ln\lambda_0+\frac{\sigma_z^2}{2\lambda_0}D-\frac{\sigma_z^2}{2\lambda_0^2}B.
\end{aligned}$$

The last approximation is made assuming the terms $\frac{\sigma_z^4}{8\lambda_0}D^2$ and $\frac{\sigma_z^4}{4\lambda_0^2}C$ are both approximately zero. In our calculations we include these terms as well.

### S4 - Simulations to test accuracy of the decomposition - comparison of different environmental variance levels $\sigma_z^2$ and environmental strength $\beta_z$

**Simulation framework:**

We used simulations to test the accuracy of the decomposition in Supporting information S3. For each of the 13 scenarios (described in the main text, Fig. 1) and for each selected population, 4,000 simulated time series of population growth were generated, each representing 200 years. For each simulated year a value of the environment $z$ was drawn from a normal distribution with mean E[$z$]=0 and variance Var($z$) $= \sigma_z^2$ (IID among time steps), and used to calculate the corresponding projection matrix, resulting in a time series of projection matrices.

Starting from a population of 100 individuals, equally distributed among the (st)ages, we computed the stochastic population growth rate for each of the 4000 simulations as the average growth increment of the total reproductive value of the population ($V_t = \sum_{j=1}^{k} v_j n_{t,j}$, where $v_j$ is the reproductive value of stage $j$ and $n_{t,j}$ is the size of (st)age $j$ at time $t$), using the estimator $ln(\lambda_s) = \frac{1}{200}\sum_{t=1}^{200} lnV_t - lnV_{t-1}$ (Engen *et al.*, 2009). The reproductive value vector $\mathbf{v}$ and the stable stage structure $\mathbf{u}$ were calculated from the projection matrix of the mean environment A(0), and $\mathbf{v}$ scaled so that $\mathbf{vu} = 1$. We then took the average $ln(\lambda_s)$ value from the 4000 simulations as the estimate for the given population and scenario. The reproductive values act as a filter for fluctuations in the (st)age structure, removing bias due to the populations being further away from the stable structure in the beginning of simulations (Engen *et al.*, 2009).

Estimating the stochastic population growth rate $ln(\lambda_s)$ using V the total reproductive value of the population, or N the population size ($ln(\lambda_s) = \frac{1}{200}\sum_{t=1}^{200} lnN_t - lnN_{t-1}$), yielded similar results (Fig. S4; Mean squared errors MSE $<$ 1.503e-05 and $R^2 \in [0.624 - 0.977]$ in the four scenarios without bony fish MPMs; MSE $<$ 6.036e-05 and $R^2 \in [0.383 - 0.995]$ with bony fishes). The greatest difference was observed for bony fish MPMs. However, the variance-covariance was affected and reduced using $V_t$. As shown by Engen *et al.* (2009), if $ln(\lambda_s)$ is calculated from the dynamics of $N_t$ the variance of annual population growth rates will be affected by autocorrelation in N generated by transient fluctuations in stage structure, typically the variance of $V_t$ is lower than that of $N_t$.

See R code for stochastic perturbations in Supporting information S7.

**Estimation of nonlinearity and variance-covariance effects on $ln(\lambda_s)$ using simulations:**

The 'small-noise' approximation of Tuljapurkar (1982) for an environment with no autocorre-

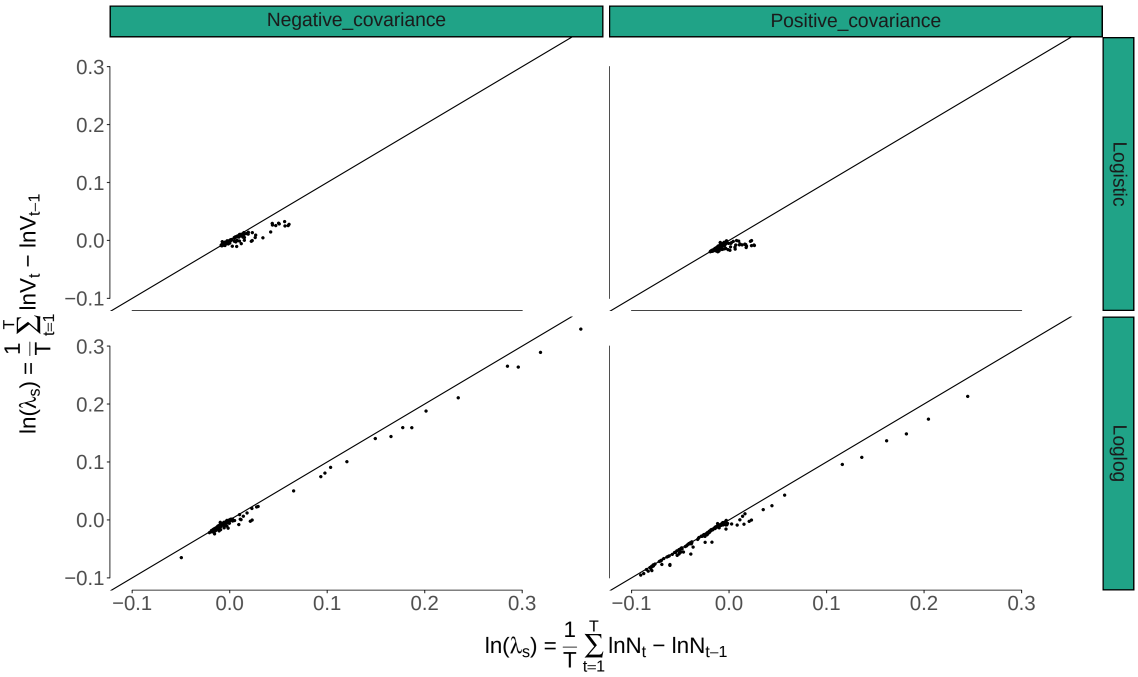


Figure S4: Comparison of the stochastic population growth rate estimated from the dynamics of the total reproductive value $V_t$ (y-axis) or of the population size $N_t$ (x-axis) under four scenarios of covariance and link-functions of the demographic parameters.

lation is given by:

$$ln(\lambda_s) \approx ln(\lambda_0) - \frac{\sigma_\lambda^2}{2\lambda_0^2} \tag{6}$$

Here, $\lambda_0$ is the dominant eigenvalue of the matrix of the mean environment $\mathbf{A(0)}$, and $\sigma_\lambda^2$ is the variance of the annual population growth rate caused by temporal variance and covariance in the matrix elements (vital rates in age structured populations). Since all the MPMs from COMADRE were rescaled so that $\lambda_0 = 1$, $\ln(\lambda_0) = 0$ in our framework.

Including nonlinear effects, the stochastic growth rate could be approximated as:

$$ln(\lambda_s) \approx ln(\lambda_0) + (ln(\bar{\lambda}) - ln(\lambda_0)) - \frac{\sigma_\lambda^2}{2\lambda_0^2} \tag{7}$$

with $\bar{\lambda}$ the dominant eigenvalue of the average matrix across variable environments $\bar{\mathbf{A}}$. Under the assumption of small fluctuations in the demographic parameters (i.e., weak and linear relationships with environment), $ln(\bar{\lambda})$ would equal $ln(\lambda_0)$. However, in case of nonlinear demographic parameter responses to environmental variability, not only the variance, but also the mean of demographic parameters can change and thereby change the arithmetic mean $\bar{\lambda}$.
For a given life history, the difference $ln(\bar{\lambda})$ - $ln(\lambda_0)$ generated by nonlinear effects of environmental variability on the demographic parameters, is a measure of overall nonlinearity effects,

and $\sigma^2 = \frac{\sigma^2_\lambda}{2\lambda_0^2}$ is a measure of the variance-covariance effects on $ln(\lambda_s)$. A positive $ln(\bar{\lambda})$ - $ln(\lambda_0)$ indicates that the overall effect of lability in demographic parameters is positive, and requires that some demographic parameters show convex relationships with the environmental driver. The value of $\sigma^2$ reflects the reduction in fitness due to variance and covariance of the demographic parameters, and can be reduced through buffering.

Using the simulation framework, we tested the accuracy of the approximation by comparing $ln(\lambda_s) = \frac{1}{200}\sum_{t=1}^{200} lnV_t - lnV_{t-1}$ (average value from the 4000 simulations) with the right side of Eq. 7 ($ln(\lambda_0) + (ln(\bar{\lambda}) - ln(\lambda_0)) - \frac{\sigma^2_\lambda}{2\lambda_0^2}$; see Supporting information S7 for calculations in R). Although simulations in our study were not limited to small fluctuations in the demographic parameters, Equation (7) provided an accurate approximation of the stochastic growth rate (Mean squared errors MSE < 3.7e-06 and $R^2 > 0.991$ in all scenarios; Fig. S5 with $N_{MPMs}$ = 129, bony fish populations are not shown). Note that the approximation is less accurate for scenarios including some demographic parameters as loglog link functions of z (i.e harsh relationships) with positive covariation, especially for bony fish populations for which the MPMs from COMADRE had very high fertility coefficients and low mean survival rates ($R^2 = 0.678$ and MSE = 6.2e-04 when bony fishes are included, instead of $R^2 = 0.994$ and MSE =3.7e-06 without those populations; see more details in SI S6).

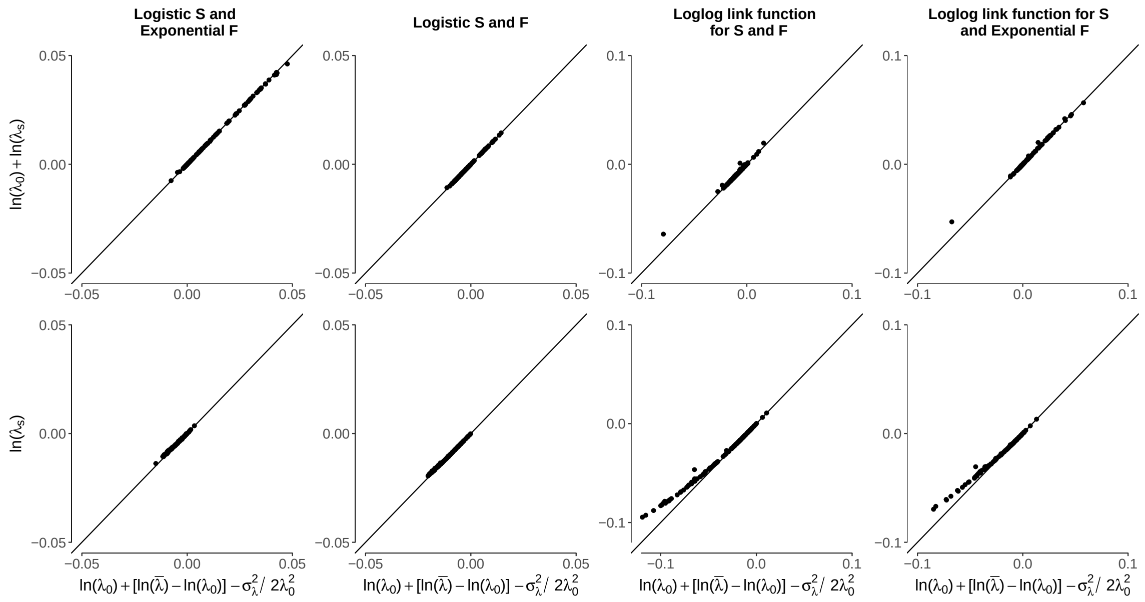


Figure S5: Tuljapurkar's approximation was found accurate for scenarios 1-8 (Fig 1 in the main part of the manuscript) with $\beta_{z_F}$ and $\beta_{z_S}$ equal to 0.4 and $\sigma^2_z$ equals to 1. Negative (upper panel) and positive (lower panel) covariation between (st)age-specific survival rates and fertility coefficients were applied. $N_{MPMs}$ = 129, results for bony fish populations are not shown.

**Nonlinearity index (D):**

To assess the net impact of demographic parameter curvatures on the direction and strength of the mean response $\ln(\bar{\lambda})$, the nonlinearity index was defined (equation 2 in the main text of the manuscript) as

$$\mathrm{D} = \sum_{ij} S_{ij} A''_{ij}(0),$$

where the sum is over all stages. The matrix $\mathbf{S}$ is the standard sensitivity matrix defined for $\mathbf{A}(0)$, with elements $\frac{d\lambda_0}{dA_{ij}} = v_i u_j$. The reproductive value vector $\mathbf{v}$ and the stable stage structure $\mathbf{u}$ are also calculated for this matrix, and $\mathbf{v}$ scaled so that $\mathbf{vu} = 1$. If an element (here demographic parameter) of the second derivative matrix is positive, the demographic parameter function is convex at $z = 0$. The sensitivities measure the influence of each matrix element on $\lambda_0$ in the mean environment, and are used to weight the impact of the curvatures to estimate the total effect of nonlinearity. Thus, a demographic parameter with strong convex curvature may still have a low impact on D if the corresponding sensitivity of $\lambda_0$ to that demographic parameter is low. The Nonlinearity index $D$ captures the overall effect of nonlinearity (i.e, strongly correlated with the difference $ln(\bar{\lambda})$ - $ln(\lambda_0)$) and indicates whether it is positive (adaptive) or negative (not adaptive). D was estimated for each population in the 13 scenarios for $\beta_{z_{0.4}}$.

**Testing for different $\sigma_z^2$ and $\beta_z$ values:**

In our simulation and analytical frameworks (main text of the manuscript), $\beta_{z_F}$ and $\beta_{z_S}$ were set to 0.4 and the environmental variance $\sigma_z^2$ was fixed to 1. To ensure that the simulated environmental variance and environmental impact on $s_j(z)$ and $f_j(z)$ were representative of a broad range of environmental variability that cause gradual changes in the response variable $\ln(\lambda_s)$ and its nonlinearity and variance-covariance components, we performed additional scenarios. Twenty scenarios were applied, combining four $\beta_z$ values ($\beta_z$ = 0.2, 0.3, 0.4 or 0.5) and five $\sigma_z^2$ values ($\sigma_z^2$ = 0.1, 0.5, 1, 1.5 or 2; Fig.S5). In all models, we assumed logistic responses for $s_j(z)$ and $f_j(z)$ (M = 2.5) to the main environmental driver z and negative covariation between them.

Our results showed that altering the $\sigma_z^2$ and/or $\beta_z$ values only affects the magnitude of the effects. Increasing the coefficients strengthens the positive and negative effects, with gradual changes. Results for the stochastic growth rate $\ln(\lambda_s)$ and its main components across generation are shown in Fig. S7-S8, respectively ($N_{MPMs}$ = 129, bony fish populations and populations with generation time $>$ 62 years are not shown).

**Comparison of methods:**

We concluded that the analytical approximation and simulation methods for estimating $\ln(\lambda_s)$ and the nonlinearity and variance-covariance components yield the same results (Spearman's rank correlation coefficients $> 0.99$ between the two approaches across all MPMs and for all scenarios, see Table 1; Fig. S8-S9). This result validates the accuracy of the decomposition of the stochastic growth rate including nonlinear effects (equation 3 in main text; Supplementary Information S3).

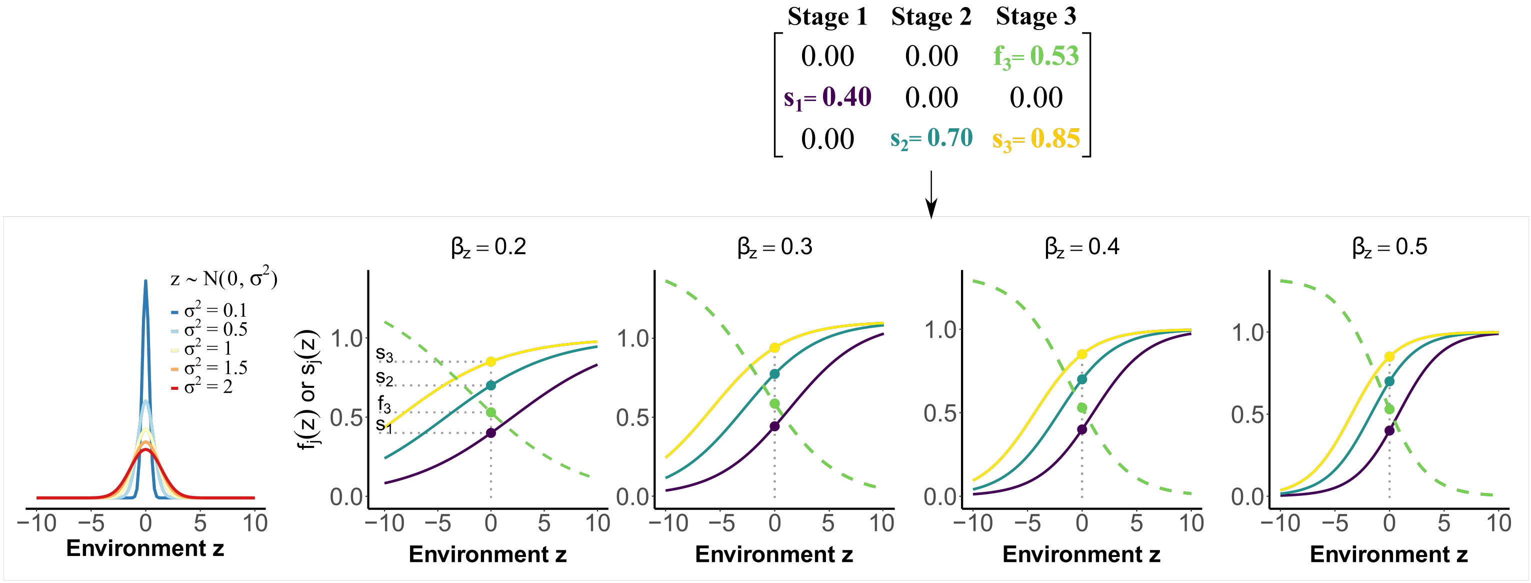


Figure S6: Different slopes $\beta_z$ (0.2-0.5) and environmental variability levels $\sigma_z^2$ (0.1-2) were investigated under 20 scenarios and illustrated here based on one matrix population model. Logistic functions (M= 2.5) and negative covariation between survival probabilities $s_j$ and fertility coefficients $f_j$ were considered. Demographic outcomes are shown on Fig. S7-S8 (analytical approach) and Fig. S9 (simulation approach).

| Spearman correlation coefficients | | | | |
|---|---|---|---|---|
| | $\beta_z = 0.2$ | $\beta_z = 0.3$ | $\beta_z = 0.4$ | $\beta_z = 0.5$ |
| 0.1 | 0.993, 0.997, 0.996 | 0.995, 0.998, 0.996 | 0.996, 0.999, 0.996 | 0.996, 0.999, 0.995 |
| 0.5 | 0.996, 0.999, 0.996 | 0.996, 0.999, 0.995 | 0.996, 0.999, 0.995 | 0.996, 0.999, 0.994 |
| $\sigma_z^2 = 1$ | 0.996, 0.999, 0.995 | 0.996, 1.000, 0.995 | 0.996, 0.999, 0.994 | 0.995, 0.999, 0.992 |
| 1.5 | 0.997, 1.000, 0.995 | 0.996, 0.999, 0.994 | 0.995, 0.999, 0.993 | 0.994, 0.998, 0.989 |
| 2 | 0.996, 1.000, 0.995 | 0.996, 0.999, 0.994 | 0.995, 0.999, 0.990 | 0.993, 0.997, 0.984 |

Table 1: Spearman correlation coefficients between the simulation and analytical approaches for three demographic outcomes estimated across all MPMs: $ln(\lambda_s)$ (black) and its nonlinearity (red) and variance-covariance (blue) components. For each MPM, survival and fertility covaried negatively and were logistic link functions of the environment z. Different slopes $\beta_z$ and environmental variances $\sigma_z^2$ were considered (Fig. S6-S9).

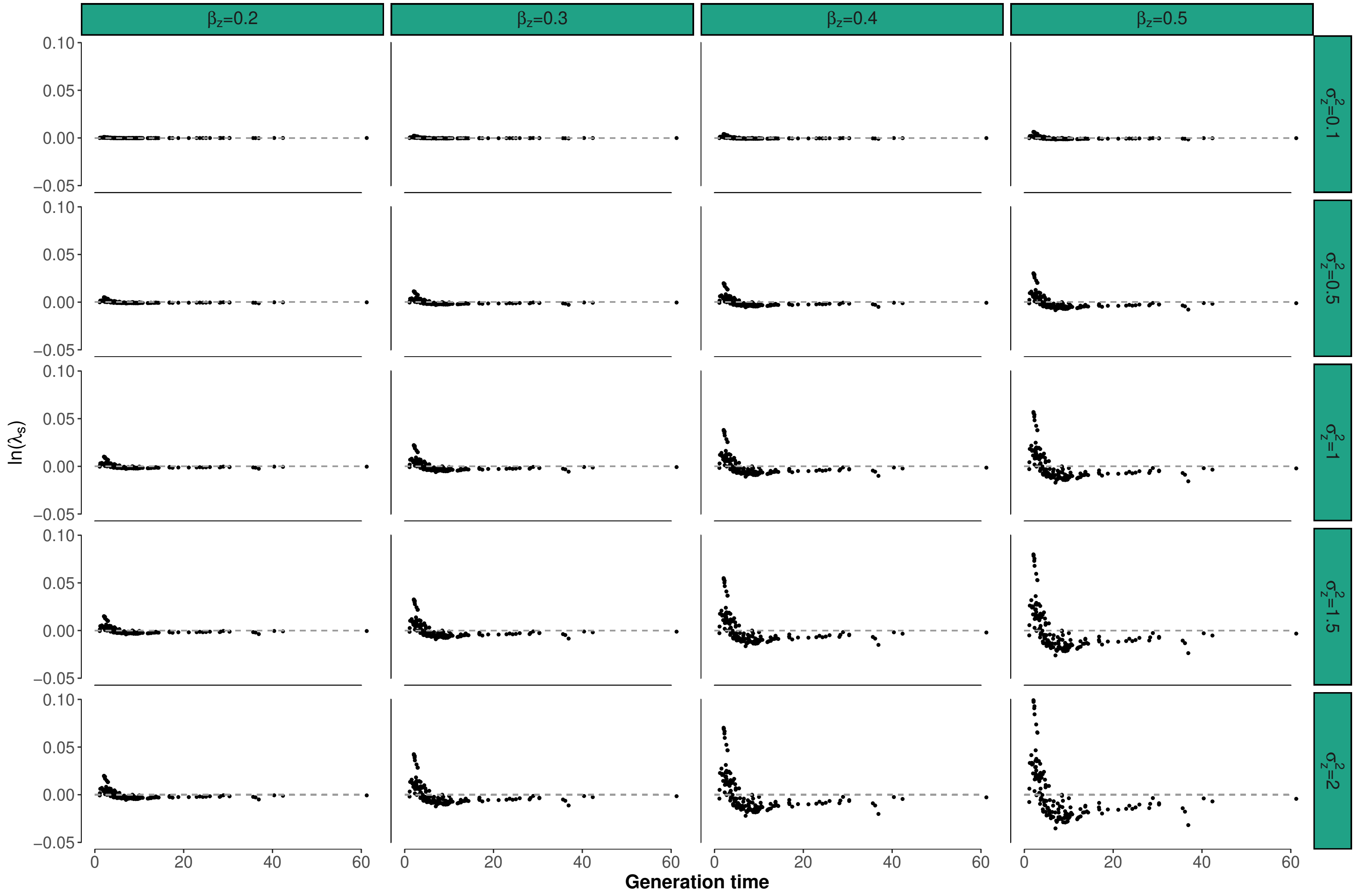


Figure S7: Analytical approach: Relationship between $ln(\lambda_s)$ and generation time when survival probabilities $s_j$ and fertility coefficients $f_j$ covary negatively and are logistic functions of the environment z (M = 2.5), using different slopes $\beta_z$ and environmental variability levels $\sigma_z^2$.

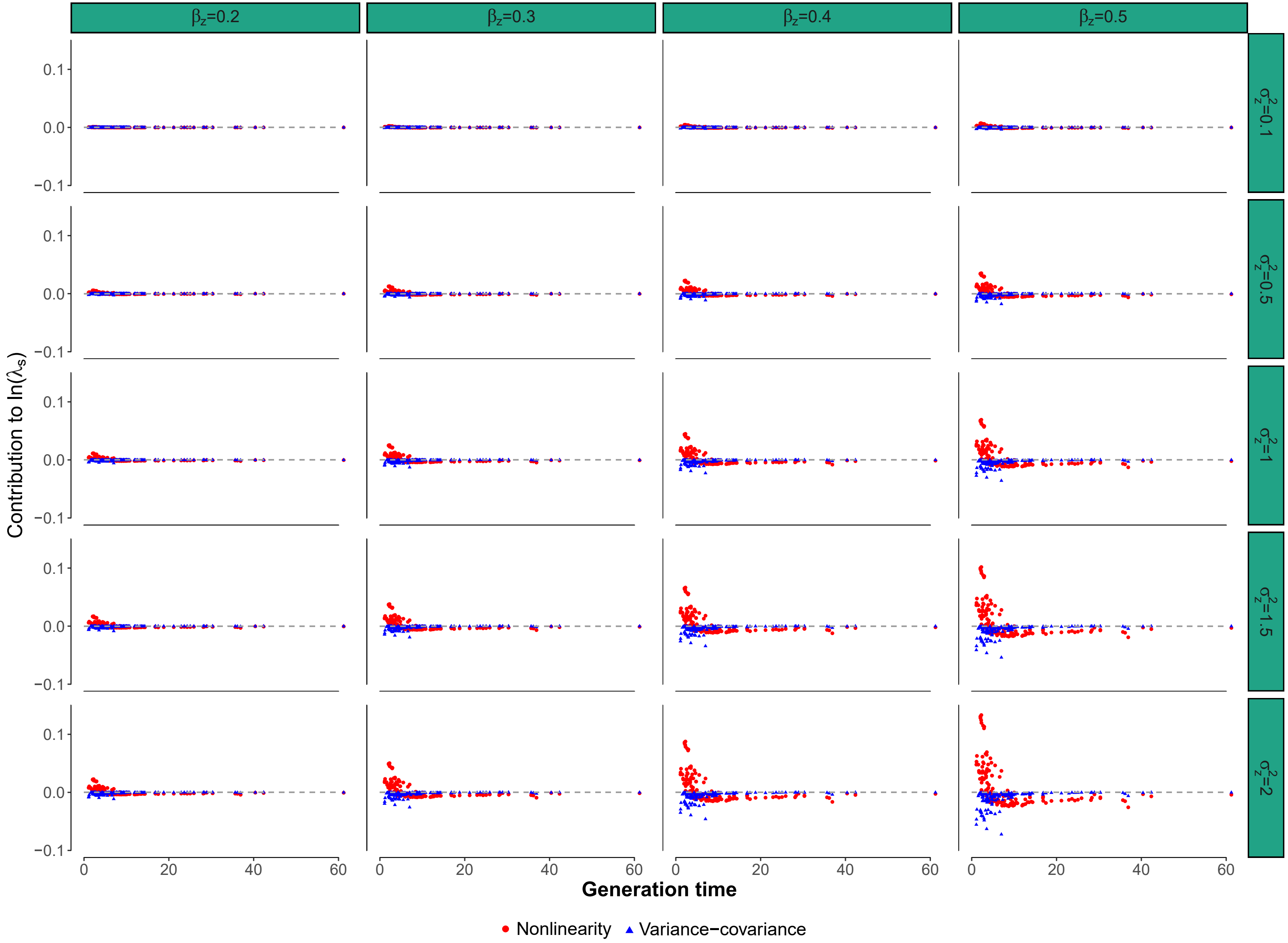


Figure S8: Analytical approach: contribution of nonlinearity (red) and variance-covariance (blue) of demographic parameters to the stochastic population growth rate $ln(\lambda_s)$, when survival probabilities $s_j$ and fertility coefficients $f_j$ covary negatively and are logistic functions of the environment z (M = 2.5), using different combinations of slopes $\beta_z$ and environmental variances $\sigma_z^2$.

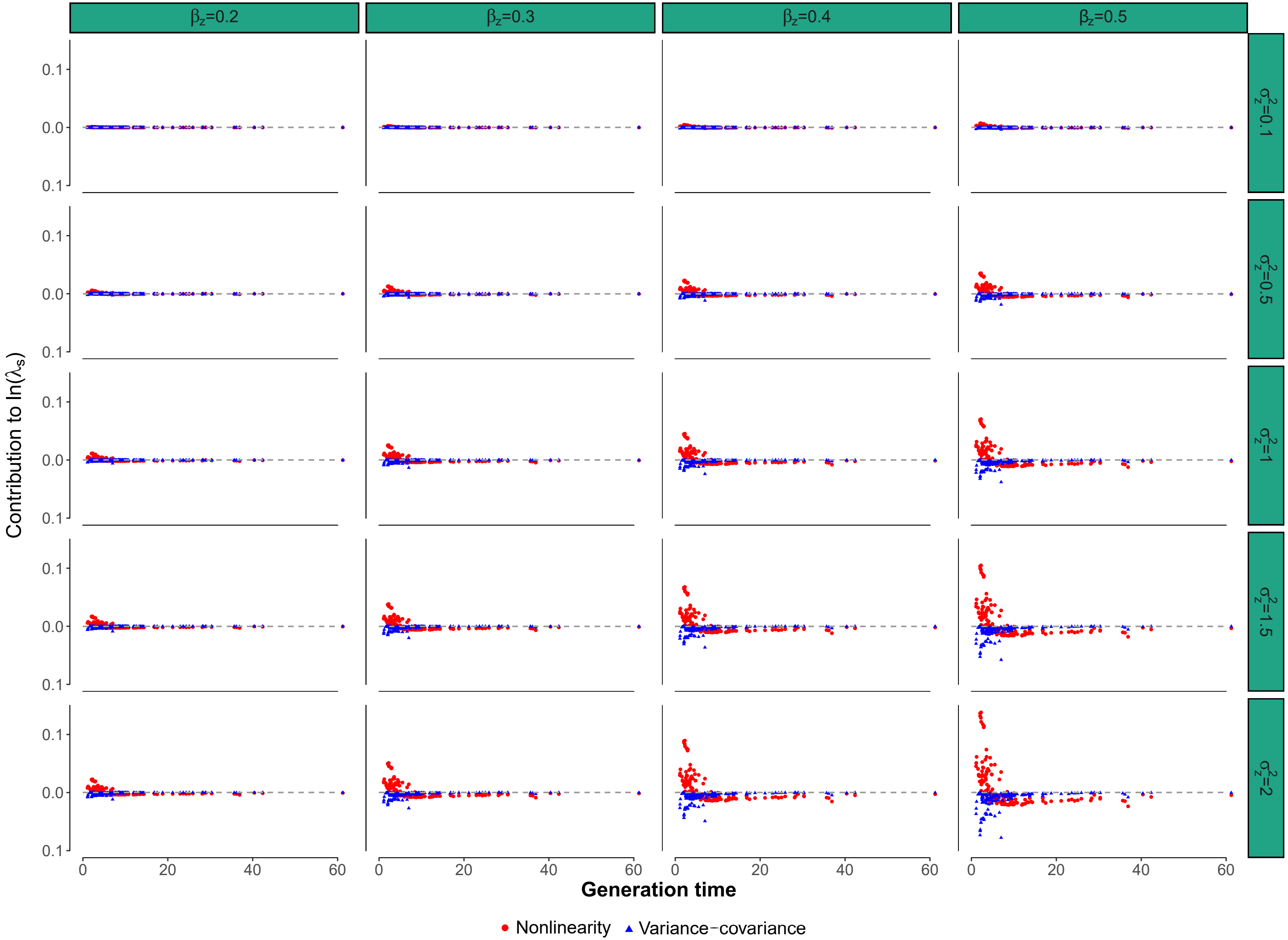


Figure S9: Simulation approach: contribution of nonlinearity (curvature, in red) and variance-covariance (in blue) of the demographic parameters to the stochastic population growth rate $ln(\lambda_s)$, when survival probabilities $s_j$ and fertility coefficients $f_j$ covary negatively and are logistic functions of the environment z (M = 2.5), using different combinations of slopes $\beta_z$ and environmental variances $\sigma_z^2$.

### S5 - Comparison of threshold values in (st)age-specific fertility coefficients

By definition, survival rates ranged between 0 and 1, while the range of (st)age-specific fertility coefficients $f_j(z)$ can vary between 0 and any positive and biologically relevant value. We used a proportional threshold where the average fertility coefficient of a (st)age j was limited to $f_j(z)$ $\in$ [0, M * $f_j(0)$] with M = 2.5 (see results in the main part of the manuscript) and $f_j(0)$ the average fertility coefficient of (st)age j reported in the COMADRE projection matrix.

In this supporting information, we compared the effects of environmental variability on the stochastic population growth rate and its main components when the proportional threshold for fertility coefficients was fixed to a lower or higher value using M = 1.5 and 3. The three different thresholds are illustrated in Fig. S10 and results are shown in Fig. S11-S12, when survival and fertility were logistic or loglog link functions of the environment z and covary positively or negatively, respectively. Our results show that populations respond in a qualitatively similar manner to environmental variability and that the magnitude of the responses increased gradually with increasing thresholds.

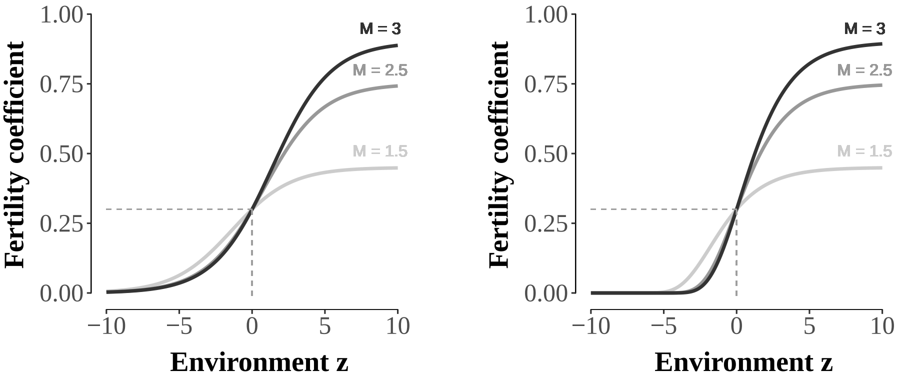


Figure S10: Fertility coefficients as logistic function (left plot) and loglog link function (right plot) of the environment z, with $\beta_{z_F}$= 0.5 (high environmental variability scenario). Following equations 1 and 2 (Supp. Information 2), MaxF (maximum fertility coefficient)= M * $f_j(0)$ with M= 1.5, 2.5 or 3 (light grey, dark grey and black lines, respectively), limiting the range of (st)age-specific fertility coefficients to $f_j(z) \in$ [0, M * $f_j(0)$]. In this example, $f_j(0)$ equals 0.27.

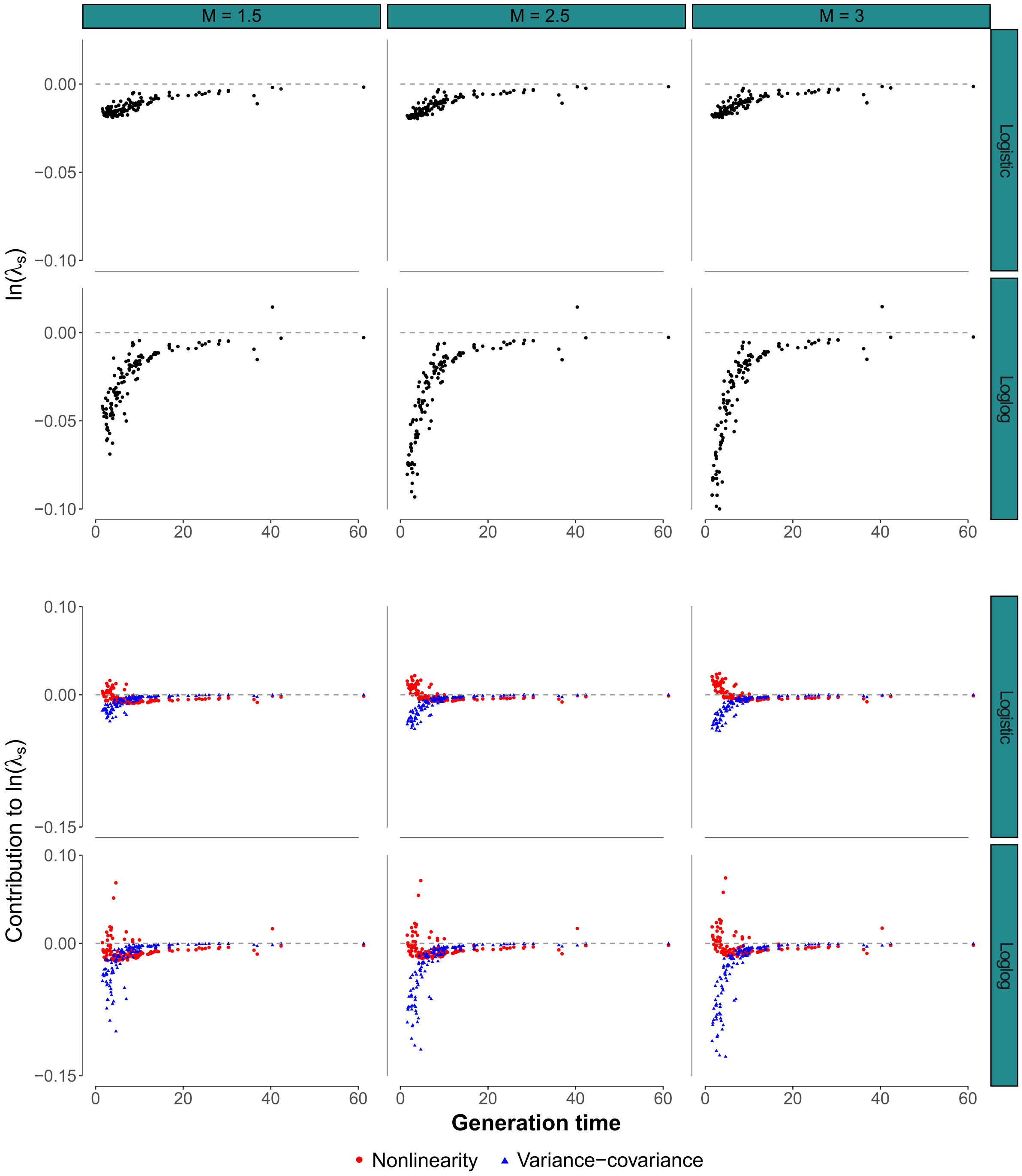


Figure S11: Effects of environmental variability on $ln(\lambda_s)$ (top panel) and on the contribution of nonlinearity and variance-covariance to $ln(\lambda_s)$ (bottom panel) across generation time. Here, survival probabilities $s_j$ and fertility coefficients $f_j$ are considered as logistic or loglog link functions of the environment z (with $|\beta_z| = 0.4$, $\sigma_z^2 = 1$) with a positive covariance between them. M = 1.5, 2.5 or 3, limiting the range of (st)age-specific fertility coefficients to $f_j(z) \in$ [0, M * $f_j(0)$]. Each point corresponds to a population.

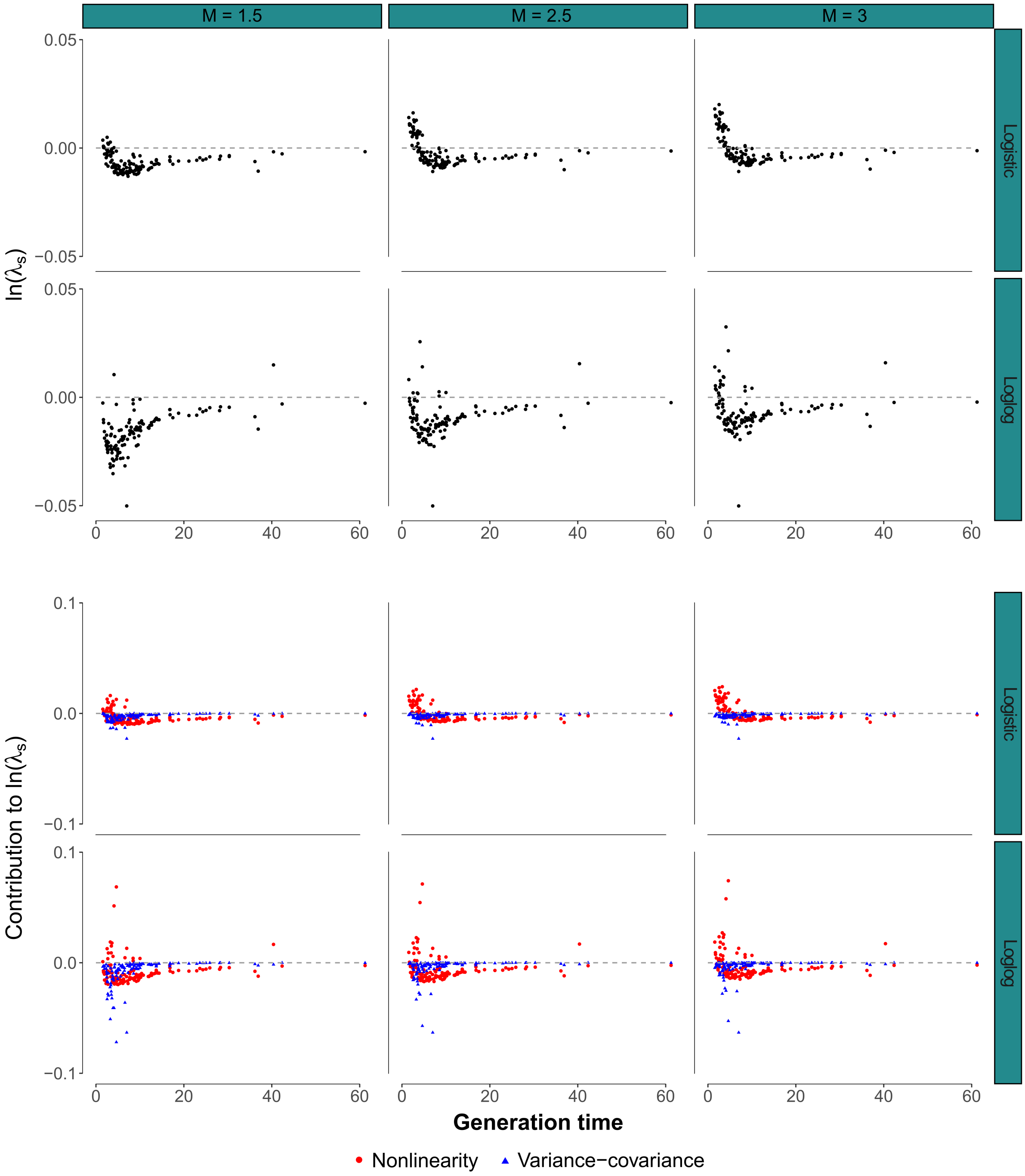


Figure S12: Effects of environmental variability on $ln(\lambda_s)$ (top panel) and on the contribution of nonlinearity and variance-covariance to $ln(\lambda_s)$ (bottom panel) across generation time. Here, survival probabilities $s_j$ and fertility coefficients $f_j$ are logistic or loglog link functions of environment z (with$|\beta_z| = 0.4$, $\sigma_z^2 = 1$) and covary negatively. M = 1.5, 2.5 or 3, limiting the range of (st)age-specific fertility coefficients to $f_j(z) \in$ [0, M * $f_j(0)$].

### S6 - Results including matrix population models of bony fishes

Nearly all matrix population models of bony fish species ($N_{MPMs}$ = 22) include very high fertility coefficients and some very low survival rates for immature stages in particular. The average fertility coefficient (offspring produced to next year) in these populations is 750, ranging from 0.9 for a population of Red groupers (*Epinephelus morio*) to 3344.9 for a population of Lake chubsuckers (*Erimyzon sucetta*). The combination of low survival rates and high fertility coefficients led to very high variance-covariance effects and thus, very low $ln(\lambda_s)$, especially under scenarios with loglog link functions for some demographic parameters. Results are shown below (all MPMs are shown, except three populations with generation time > 62 years, $N_{MPMs}$ = 151; Fig. S14-16).

In Supporting information S4, we showed that the Tuljapurkar's approximation of the stochastic growth rate including nonlinear effects was robust to small to moderate fluctuations in demographic parameters for our selected MPMs. Under scenarios of loglog link functions for survival rates and/or fertility coefficients (Fig. S14b-d and S15b-d), this assumption was violated for most of the bony fish populations due to the extreme mean and variance of survival rates and fertility coefficients. The highest the variance, the lowest the accuracy of the variance-covariance component using our calculation and decomposition (Fig. S13). Scenarios should be adjusted using more specific and biologically realistic assumptions for these life histories (e.g using smaller amplitudes of variation for the fertility coefficients or the survival rates of immature stages.

For other scenarios using logistic functions (Fig. S14a-c, S15a-c and S16), although the magnitude of responses is higher for these species, the trend and interpretation of the results remain the same as those presented in the main body of the manuscript (Fig. 2-3).

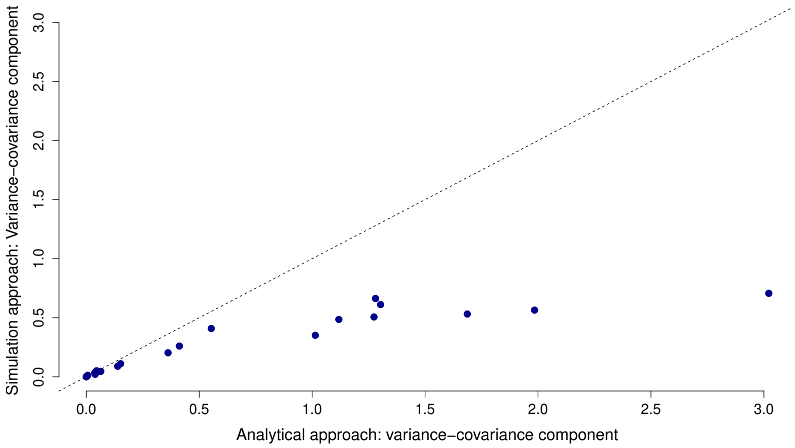


Figure S13: Comparison between the simulation and analytical approaches of the variance-component estimated for the bony fish MPMs ($N_{MPMs} = 22$). The approximation is accurate for small fluctuations in demographic parameters but became unrealistic as the variance increased.

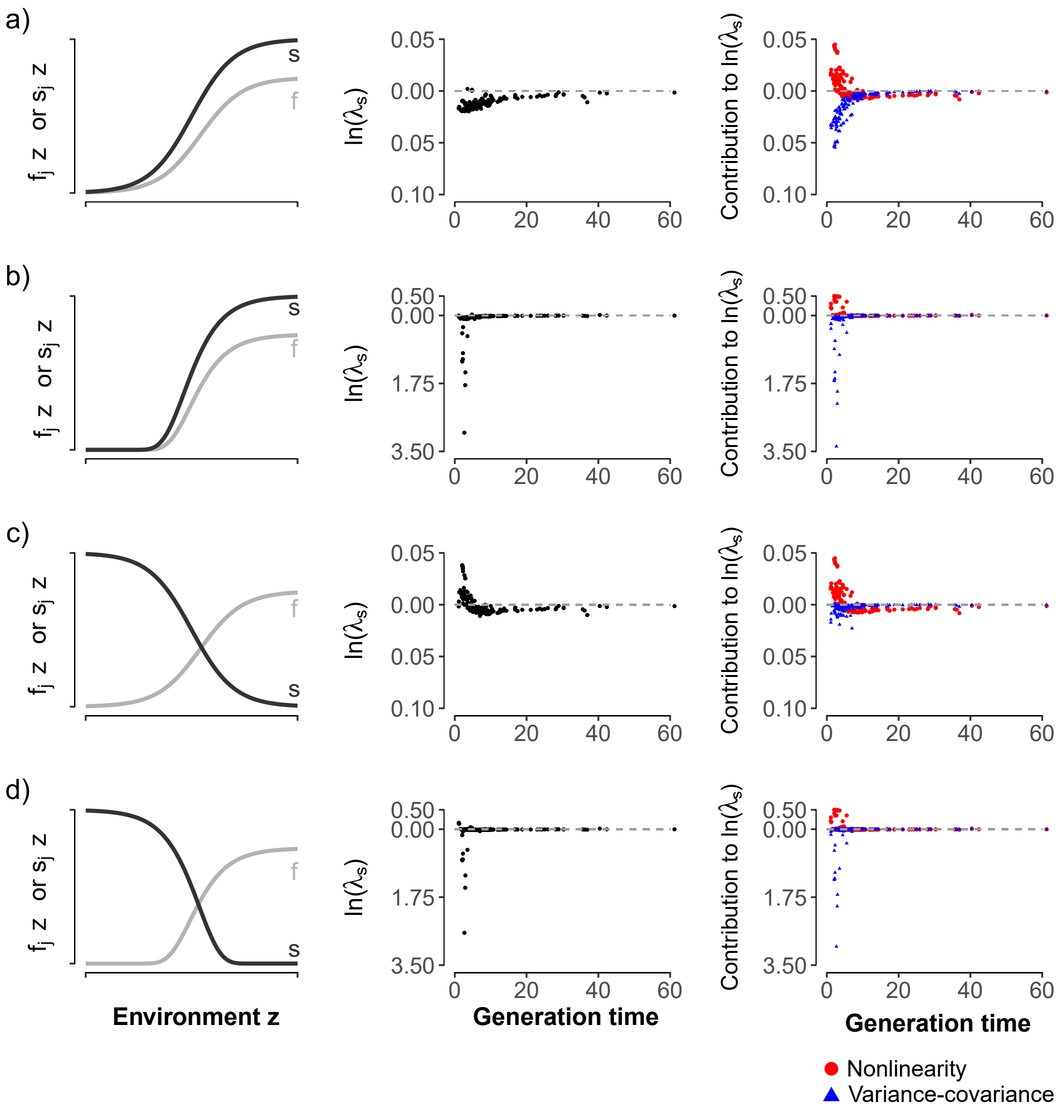


Figure S14: Mid panels: Effects of environmental variability on the long-term fitness $ln(\lambda_s)$ across generation time, under four scenarios of covariance and link-functions of the demographic parameters. The four scenarios are illustrated on the left panel, with the grey and black lines corresponding to the (st)age-specific survival rates $s_j(z)$ and fertility coefficients $f_j(z)$, respectively (functions varied for each stage depending on $s_j(0)$ and $f_j(0)$; only one function is shown for survival and fertility here). For each population, positive (panels a-c) and negative (panels b-d) covariance between $f_j(z)$ and $s_j(z)$ were simulated, considering $f_j(z)$ and $s_j(z)$ as logistic functions (panels a-b) or loglog link functions (panels c-d) of the environment z ($|\beta_z| = 0.4$, $\sigma_z^2 = 1$). Panels to the right show the decomposition of $\ln(\lambda_s)$ into main components capturing variance-covariance effects (blue triangles) and nonlinear effects on the mean growth rate $\ln \bar{\lambda}$ (lability, red circles).

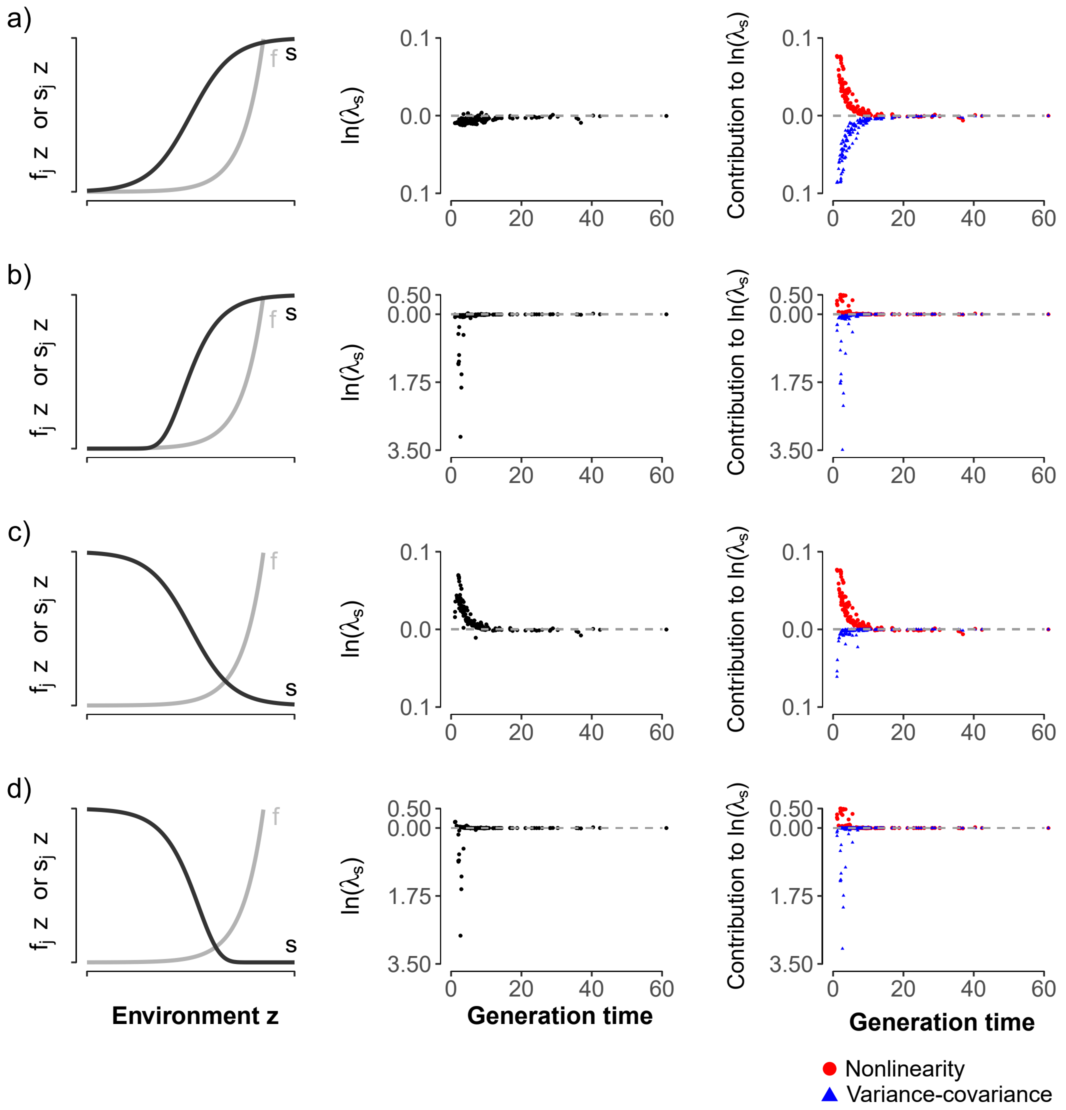


Figure S15: Mid panels: Effects of environmental variability on the long-term fitness $ln(\lambda_s)$ across generation time, using logistic (panels a-b) or loglog (panels c-d) link functions for (st)age-specific survival rates $s_j(z)$, and log link function for fertilities $f_j(z)$ ($|\beta_z| = 0.4$, $\sigma_z^2 = 1$). For each population, positive (panels a-c) and negative (panels b-d) covariance between (st)age-specific survival rates and fertilities were simulated. Panels to the right show the decomposition of $ln(\lambda_s)$ into main components capturing variance-covariance effects (blue triangles) and non-linear effects on the mean growth rate $\ln \bar{\lambda}$ (lability, red circles).

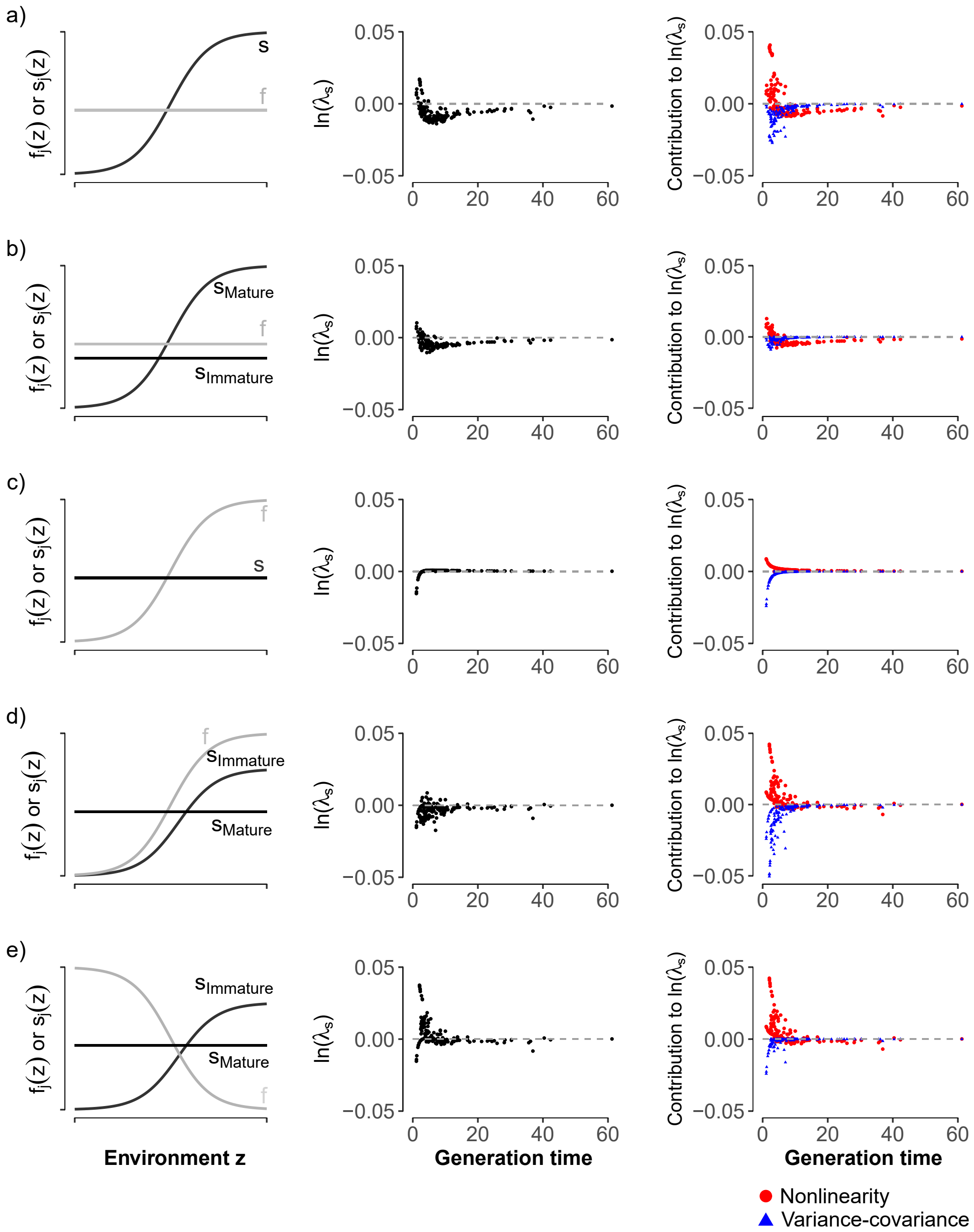


Figure S16: Mid panels: Results from scenarios of forced buffering with demographic lability only among (a) (st)age-specific survival rates, (b) survival rates of the reproductive stages only, (c) (st)age-specific fertilities and (d-e) fertilities and survival rates of the non-reproductive stages. For each scenario, the long term fitness $\ln(\lambda_s)$ and its main components reflecting nonlinearity (red circles) and variance covariance (blue triangles) effects are plotted against generation time (mid and right panels).

### S7 - R code: decomposition of the stochastic population growth rate

The code below shows how to decompose the stochastic population growth into the nonlinearity and variance-covariance components, using both the analytical and simulation approaches. An example is given at the end of the code, considering (st)age-specific survival probabilities $s_j(z)$ and fertility coefficients $f_j(z)$ for each population as logistic functions of the environmental variable z, with mean E$[z]$=0 and variance $\sigma_z^2$= 1. In all simulations, we considered that the steepness and direction of the relationship between z and the survival rates or the fertility coefficients, were the same for all age classes or stages in a population. In other words, we assumed that survival rates of different (st)ages have the same value of $\beta_{z_S}$, and similarly all fertility coefficients have the same $\beta_{z_F}$ ($|\beta_{z_F}| = |\beta_{z_S}| = 0.4$).

```
1. First and second derivatives, and functions to be used later
=================================================================================
Defining the first (fdev) and second (sdev) derivatives of the functions relating
the (st)age-specific fertility and survival rates to the environmental variable z:
=================================================================================
#Inputs:
#z: the environmental vector
#Bm: fertility coefficient of (st)age j in the COMADRE matrix (at z=0)
#Meff = 1.5, 2.5 or 3 - different thresholds for fertility coefficients, with f_j(z)
    in [0, Meff * f_j(0)]
#fert.eff or surv.eff = 0.5 or 0.1 to simulate high or low environmental variability
    effects on demographic parameters, respectively.

****************
#Logit function:
****************
logit<-function(x){
  log(x/(1-x))
}
***************************************************
#Fertility coefficients as logistic functions of z:
***************************************************
#Function
fertility.logistic <- function(B= Bm, z, bz= fert.eff, maxeff= Meff){
  maxF <- maxeff * B # Maximum fertility
```

```
  int.mu <- logit(1/maxeff)
  maxF/(1+exp(-int.mu-bz*z))
}
#First derivative
fdev.fertility.logistic <- function(B= Bm, z, bz= fert.eff, maxeff= Meff){
  maxF <- maxeff * B
  int.mu <- logit(1/maxeff)
  A <- exp(-(int.mu+bz*z))
  maxF*bz*A/((1+A)^2)
}
#Second derivative
sdev.fertility.logistic <- function(B= Bm, z, bz= fert.eff, maxeff= Meff){
  maxF <- maxeff *B
  int.mu <- logit(1/maxeff)
  A <- exp(-(int.mu+bz*z))
  2* maxF*bz^2*A^2/(1+A)^3- maxF*bz^2*A/(1+A)^2
}

*****************************************************
#Fertility coefficients as loglog link functions of z:
*****************************************************
#Function
fertility.loglog <- function(B= Bm, z, bz= fert.eff, maxeff= Meff){
  maxF <- maxeff * B # Maximum fertility
  int.mu <- -log(-log(1/maxeff)) # intercept of linear contrast loglog link
  maxF*exp(-exp(-int.mu-bz*z))
}
#First derivative
fdev.fertility.loglog <- function(B= Bm, z, bz= fert.eff, maxeff= Meff){
  maxF<- maxeff * B
  int.mu<- -log(-log(1/maxeff))
  maxF*bz*exp(-int.mu-bz*z)*exp(-exp(-int.mu-bz*z))
}
#Second derivative
sdev.fertility.loglog <- function(B= Bm, z, bz= fert.eff, maxeff= Meff){
  maxF<- maxeff * B
  int.mu <- -log(-log(1/maxeff))
  maxF* bz^2*exp(-int.mu-bz*z)*exp(-exp(-int.mu-bz*z))*(exp(-int.mu-bz*z)-1)
}
```

```
**************************************************
#Fertility coefficients as log link functions of z:
**************************************************
#Function
fertility.log <-function(B = Bm, z, bz= fert.eff){
  int.b <- log(B) # intercept of linear contrast log link
  exp(int.b+bz*z)
}
#First derivative
fdev.fertility.log <- function(B= Bm, z, bz= fert.eff){
  int.b <- log(B)
  bz*exp(int.b+bz*z)
}
#Second derivative
sdev.fertility.log <- function(B= Bm, z, bz= fert.eff){
  int.b <- log(B)
  bz^2*exp(int.b+bz*z)
}

******************************************
#Survival rates as logistic functions of z:
******************************************
#Function
survival.logistic <- function(S= Sm, z= 0, bz= surv.eff){
  int.mu <- logit(S)
  1/(1+exp(-int.mu-bz*z))
}
#First derivative
fdev.survival.logistic <- function(S= Sm, z= 0, bz= surv.eff){
  int.mu <- logit(S)
  A <- exp(-(int.mu+bz*z))
  bz*A/((1+A)^2)
}
#Second derivative
sdev.survival.logistic <- function(S= Sm, z= 0, bz= surv.eff){
  int.mu <- logit(S)
  A <- exp(-(int.mu+bz*z))
  2*bz^2*A^2/(1+A)^3- bz^2*A/(1+A)^2
}
```

```
**********************************************
#Survival rates as loglog link functions of z:
**********************************************
#Function
survival.loglog <- function(S= Sm, z= 0, bz= surv.eff){
  int.mu <- -log(-log(S)) # intercept of linear contrast loglog link
  exp(-exp(-int.mu-bz*z))
}
#First derivative
fdev.survival.loglog <- function(S= Sm, z= 0, bz= surv.eff){
  int.mu <- -log(-log(S))
  bz*exp(-int.mu-bz*z)*exp(-exp(-int.mu-bz*z))
}
#Second derivative
sdev.survival.loglog <- function(S= Sm, z= 0, bz= surv.eff){
  int.mu <- -log(-log(S))
  bz^2*exp(-int.mu-bz*z)*exp(-exp(-int.mu-bz*z))*(exp(-int.mu-bz*z)-1)
}

======================================================================
Functions to calculate demographic quantities from a projection matrix
======================================================================
***************************************************************************
#wvlambda returns the stable structure "w", the reproductive values "v" and
#the asymptotic growth rate "lambda" given a projection matrix
***************************************************************************
#Input: matA: projection matrix
wvlambda<-function(MatA){
  ev <- eigen(MatA)
  tev <- eigen(t(MatA))
  lmax<-which.max(Re(ev$values))
  W <- ev$vectors
  V <- tev$vectors
  w <- as.matrix(abs(Re(W[, lmax]))/sum(abs(Re(W[, lmax]))))
  v <- as.matrix(abs(Re(V[, lmax])))
  v <- v/sum(v*w)
  return(list("lambda"=max(Re(ev$values)),"w"=w,"v"=v))
}
```

```
************************************************
#MatRfunc returns the transition matrix from MatU
************************************************
#Input: matU: sub-matrix including (st)age-specific survival rates
MatRfunc <- function(MatU){
  k<-dim(MatU)[1]
  if(is.na(MatU[k,k])){
    MatU[k,k]<-0
  }
  ColsumU<-apply(MatU,2,sum)
  matR<-t(t(MatU)/ColsumU)
  for(i in 1:k){
    if(ColsumU[i]==0){
      matR[,i]<-0
      matR[i,i]<-1
    }
  }
  matR
}

**********************************************************
#MatQfunc returns the offspring transition matrix from MatF
**********************************************************
#MatF = sub-matrix including (st)age-specific fertility coefficients
MatQfunc <- function(MatF){
  k<-dim(MatF)[1]
  Qmat<-matrix(0,k,k)
  ColsumF<-apply(MatF,2,sum)
  for(i in 1:k){
    if(ColsumF[i]==0){
      Qmat[1,i]<-1
    }
    else{
      Qmat[,i]<-MatF[,i]/ColsumF[i]
    }
  }
  Qmat
}
```

```
**********************************************
#Function GenTime to calculate generation time
**********************************************
#Mean age of parents at the stable (st)age distribution (Caswell, 2001).
#Inputs:
#MatA = projection matrix
#MatF = sub-matrix including (st)age-specific fertility coefficients
GenTime <- function(MatA,MatF){
  res <- wvlambda(MatA=MatA)
  lam <- res$lam
  w <- res$w
  v <- res$v
  lam/(t(v)%*%MatF%*%w)
}

**************************************************************************************
#Function sensitivity.matrix to calculate sensitivity of lambda with respect to MatA
    elements
*************************************************************************************
#Inputs: MatA = projection matrix
sensitivity.matrix <- function(MatA, zeroes = FALSE){
  res <- wvlambda(MatA=MatA)
  w <- as.vector(res$w)
  v <- as.vector(res$v)
  sensmat <- t(w%o%v)
  if (zeroes==T){
    sensmat <- ifelse (MatA==0, 0, sensmat)
  }
  sensmat
}
```

```
*************************************************************
#Function FirstDeriv to calculate the first derivative matrix
*************************************************************
#FirstDeriv calculates the first derivatives of the (st)age-specific S and F functions
#at z=0, and recombines them into a matrix of the same dimension as the input.

#Inputs:
#MatF: sub-matrix with (st)age-specific fertility coefficients
#MatU: sub-matrix with survival and transitions among stages
#survival.link: Choice of link function for survival (logistic or loglog)
#fertility.link: Choice of link function for fertility (logistic, loglog, or log)
#betaS: Effect of z on survival on the link scale
#betaF: Effect of z on fertility on the link scale
#maxeff: Maximum fertility (if logistic or loglog)

FirstDeriv <- function(MatU, MatF, survival.link="logistic", fertility.link="logistic"
    , betaS=surv.eff, betaF=fert.eff, maxeff=2.5){
  k<-length(MatU[1,])
  MatQ <- MatQfunc(MatF)
  MatR <- MatRfunc(MatU)
  Svec <- apply(MatU,2,sum) #survival vector
  Bvec <- apply(MatF,2,sum) #fertility vector
  if(survival.link=="logistic"){
    Sdev1 <- fdev.survival.logistic(S=Svec, z=0, bz=betaS)
  }
  if(survival.link=="loglog"){
    Sdev1 <- fdev.survival.loglog(S=Svec,z=0,bz=betaS)
  }
  if(fertility.link=="logistic"){
    Bdev1 <- fdev.fertility.logistic(B=Bvec,z=0,bz=betaF, maxeff=maxeff)
  }
  if(fertility.link=="loglog"){
    Bdev1 <-fdev.fertility.loglog(B=Bvec,z=0,bz=betaF, maxeff=maxeff)
  }
  if(fertility.link=="log"){
    Bdev1 <-fdev.fertility.log(B=Bvec,z=0,bz=betaF)
  }
  t(matrix(Sdev1,k,k))*MatR+t(matrix(Bdev1,k,k))*MatQ
}
```

```
*************************************************************
#Function SecondDeriv to calculate the second derivative matrix
*************************************************************
#SecondDeriv calculates the second derivatives of the (st)age-specific S and F
#functions at z=0, and recombines them into a matrix of the same dimension as the
    input.
#Inputs:
#MatF: sub-matrix with (st)age-specific fertility coefficients
#MatU: sub-matrix with survival and transitions among stages
#survival.link: Choice of link function for survival (logistic or loglog)
#fertility.link: Choice of link function for fertility (logistic, loglog, or log)
#betaS: Effect of z on survival on the link scale
#betaF: Effect of z on fertility on the link scale
#maxeff: Maximum fertility (if logistic or loglog)

SecondDeriv <- function(MatU, MatF, survival.link="logistic", fertility.link="logistic
    ", betaS=surv.eff, betaF=fert.eff, maxeff=2.5){
  k<-length(MatU[1,])
  MatQ <- MatQfunc(MatF)
  MatR <- MatRfunc(MatU)
  Svec <- apply(MatU,2,sum) #survival vector
  Bvec <- apply(MatF,2,sum) #fertility vector
  if(survival.link=="logistic"){
    Sdev1 <- sdev.survival.logistic(S=Svec, z=0, bz=betaS)
  }
  if(survival.link=="loglog"){
    Sdev1 <- sdev.survival.loglog(S=Svec,z=0,bz=betaS)
  }
  if(fertility.link=="logistic"){
    Bdev1 <- sdev.fertility.logistic(B=Bvec,z=0,bz=betaF, maxeff=maxeff)
  }
  if(fertility.link=="loglog"){
    Bdev1 <-sdev.fertility.loglog(B=Bvec,z=0,bz=betaF, maxeff=maxeff)
  }
  if(fertility.link=="log"){
    Bdev1 <-sdev.fertility.log(B=Bvec, z=0, bz=betaF)
  }
  t(matrix(Sdev1,k,k))*MatR+t(matrix(Bdev1,k,k))*MatQ
}
```

```
=============================================================================
2. ANALYTICAL APPROACH: Tuljapurkar's approximation with non-linear effects
=============================================================================

*********************************************************
#Function Approx_rS to calculate stochastic growth rate:
*********************************************************
#Approx_rS calculates the stochastic growth rate (r_S = ln(lambda_s)) based on
#Tuljapurkar's approximation and returns this plus its main components.
#Returns a list of calculated properties
#including generation time, lambda0 and the stochastic growth rate components.

#Inputs:
#MATS: a list of matrices matA, matF and matU for a given model
#varZ: Variance of z.
#survival.link: Choice of link function for survival (logistic or loglog)
#fertility.link: Choice of link function for fertility (logistic, loglog, or log)
#betaS: Effect of z on survival on the link scale
#betaF: Effect of z on fertility on the link scale
#maxeff: Maximum fertility (if logistic or loglog)

Approx_rS <- function(MATS, varZ=1, survival.link="logistic", fertility.link="logistic
    ", betaS=surv.eff, betaF=fert.eff, maxeff= Meff){
  lambda1 <- wvlambda(MATS$matA)$lam
  MatA <- MATS$matA/lambda1 #standardization of each MPM
  MatU <- MATS$matU/lambda1
  MatF <- MATS$matF/lambda1
  lambda <-wvlambda(MatA)$lam
  if(any(apply(MatU,2,sum)>1)){
    returns <- list("rS"=NA, "NonlinearityPart"=NA, "VariancePart"=NA, "D"=NA, "B"=NA,
         "C"=NA, "lambda0"=lambda, "GenTime"=NA, "NonlinearityPartFert"=NA, "
        NonlinearityPartSurv"=NA, "lambda0.notscaled"=lambda1)
  }
  else{
    Gtime <- GenTime(MatA, MatF)
    k<-length(MatA[1,])
    Sensmat <- sensitivity.matrix(MatA,zeroes=T)
```

```
    FirstDerivMat <- FirstDeriv(MatU, MatF, survival.link = survival.link, fertility.
        link = fertility.link, betaS=betaS, betaF=betaF, maxeff=maxeff)
    SecondDerivMat <- SecondDeriv(MatU, MatF, survival.link = survival.link, fertility
        .link = fertility.link, betaS=betaS, betaF=betaF, maxeff=maxeff)
    j <- which(FirstDerivMat[]=="NaN")
    FirstDerivMat[j] <- 0
    j <- which(SecondDerivMat[]=="NaN")
    SecondDerivMat[j] <- 0
    SecondDerivFMat <- SecondDerivUMat <- matrix(0,k,k )
    SecondDerivFMat[1,] <- SecondDerivMat[1,]
    SecondDerivUMat[-1,] <- SecondDerivMat[-1,]

    #Nonlinearity index D:
    D <- sum(SecondDerivMat*Sensmat)
    DFertility<-sum(SecondDerivFMat*Sensmat) #Fertility component
    DSurvival <-sum(SecondDerivUMat*Sensmat) #Survival component
    #Linear effects on the variance (double sum):
    B <-sum(outer(FirstDerivMat*Sensmat,FirstDerivMat*Sensmat))
    #Non-linear effects on the variance(double sum)
    C <- sum(outer(SecondDerivMat*Sensmat, SecondDerivMat*Sensmat))
    #Nonlinearity component
    Nonlinearity <- varZ/(2*lambda)*D*(1-varZ/(4*lambda)*D)
    NonlinearityFert <- varZ/(2*lambda)*DFertility*(1-varZ/(4*lambda)*DFertility)
    NonlinearitySurv <- varZ/(2*lambda)*DSurvival*(1-varZ/(4*lambda)*DSurvival)
    #Variance-covariance component
    Varpart <- varZ/(2*lambda^2)*(B+varZ/2*C)
    #Total:
    rS <- log(lambda)+Nonlinearity-Varpart
    returns<-return(list("rS"=rS, "NonlinearityPart"=Nonlinearity, "VariancePart"=
        Varpart, "D"=D, "B"=B, "C"=C, "lambda0"=lambda, "GenTime"=Gtime, "
        NonlinearityPartFert"= NonlinearityFert, "NonlinearityPartSurv"=
        NonlinearitySurv, "lambda0.notscaled"=lambda1))
  }
  return(returns)
}
```

```
************************************************************************
#Function return.results to calculate results for many comadre models
************************************************************************
#For each model in a comadre subset, return.results applies the Approx_rS function to
    return the stochastic growth rate, its components, and other parameters. Returns a
     data frame of the calculated properties.

#Inputs:
#comadremodels: an object with a subset of entries from the comadre database, with the
     same structure.
#varZ: Variance of z.
#survival.link: Choice of link function for survival (logistic or loglog)
#fertility.link: Choice of link function for fertility (logistic, loglog, or log)
#betaS: Effect of z on survival on the link scale
#betaF: Effect of z on fertility on the link scale
#maxeff: Maximum fertility (if logistic or loglog)

return.results <- function(comadremodels=xdataALL, varZ=1, survival.link="logistic",
    fertility.link="logistic", betaS=surv.eff, betaF=fert.eff, maxeff= Meff){
  len <- length(comadremodels$mat)
  output <- data.frame(GenTime = rep(NA, len),
                     rS = rep(NA, len),
                     NonlinearityPart = rep(NA, len),
                     VariancePart = rep(NA, len),
                     D = rep(NA, len),
                     B = rep(NA, len),
                     C = rep(NA, len),
                     NonlinearityPartFert = rep(NA, len),
                     NonlinearityPartSurv = rep(NA, len),
                     lambda0 = rep(NA, len),
                     lambda0.notscaled = rep(NA, len))
  #2. For each comadre model
  for (i in 1:len){
    mats <- comadremodels$mat[[i]]
    #print(i)
    #Remove models with NA values in matrix
    if (any(is.na(mats$matA))|any(is.na(mats$matU))|any(is.na(mats$matF))){
      output$GenTime[i] <- NA
```

```
      output$rS[i] <- NA
      output$NonlinearityPart[i] <- NA
      output$VariancePart[i] <- NA
      output$D[i] <- NA
      output$B[i] <- NA
      output$C[i] <- NA
      output$NonlinearityPartFert[i] <-NA
      output$NonlinearityPartSurv[i] <-NA
      output$lambda0[i] <- NA
      output$lambda0.notscaled[i] <- NA
    } else {
      #Calculate results
      approx <- Approx_rS(MATS=mats, varZ=varZ, survival.link=survival.link, fertility
          .link=fertility.link, betaS=betaS, betaF=betaF, maxeff= maxeff)
      output$GenTime[i] <- approx$GenTime
      output$rS[i] <- approx$rS
      output$NonlinearityPart[i] <- approx$NonlinearityPart
      output$VariancePart[i] <- approx$VariancePart
      output$D[i] <- approx$D
      output$B[i] <- approx$B
      output$C[i] <- approx$C
      output$NonlinearityPartFert[i] <- approx$NonlinearityPartFert
      output$NonlinearityPartSurv[i] <- approx$NonlinearityPartSurv
      output$lambda0[i] <- approx$lambda0
      output$lambda0.notscaled[i] <- approx$lambda0.notscaled
    }
  }
  output
}
```

```
=====================================================
              3. SIMULATION APPROACH
=====================================================

***********************************************************
Creation of a matrix array as a function of z (vector zvec)
***********************************************************
#Inputs:
#zvec: vector of z-values (environmental variable)
#Svec: survival vector
#Fvec: fertility vector
#Rmat: transition matrix
#Qmat: offspring transition matrix
#survival.link: Choice of link function for survival (logistic or loglog)
#fertility.link: Choice of link function for fertility (logistic, loglog, or log)
#betaS: Effect of z on survival on the link scale
#betaF: Effect of z on fertility on the link scale
#maxeff: Maximum fertility (if logistic or loglog)

#Output: An array of projection matrices produced by the different z values (an
    independent and identically distributed random variable with a Gaussian
    distribution)

matrix.array <- function(zvec= z.sim, Svec, Fvec, Rmat, Qmat, survival.link="logistic"
    , fertility.link="loglog", betaS=surv.eff, betaF=fert.eff, maxeff=Meff){
  n <- length(zvec) #dimension of environment vector
  k <- length(Svec) #dimension of the projection matrix
  mat <- array(dim=c(k,k,n))
  for (i in 1:n){
    if(survival.link=="logistic"){
      Svek<-survival.logistic(S=Svec,z=zvec[i], bz=betaS)
    }
    if(survival.link=="loglog"){
      Svek<-survival.loglog(S=Svec,z=zvec[i], bz=betaS)
    }
    if(fertility.link=="logistic"){
      Bvek<-fertility.logistic(B=Fvec,z=zvec[i], bz=betaF, maxeff=maxeff)
    }
    if(fertility.link=="loglog"){
```

```
    Bvek<-fertility.loglog(B=Fvec,z=zvec[i], bz=betaF, maxeff=maxeff)
   }
   if(fertility.link=="log"){
    Bvek<-fertility.log(B=Fvec, z=zvec[i], bz=betaF)
   }

   mat[,,i]<-t(matrix(Svek,k,k))*Rmat + t(matrix(Bvek,k,k))*Qmat
  }
  mat
}

************************************************************
Projection of population size and total reproductive values
************************************************************
#Using the array of projection matrices produced by the different z values,
#the popsim function allows to project the population forward in time.

# Inputs:
# matarray: array of projection matrices (from matrix.array)
# v: Reproductive values calculated for the mean projection matrix
# N0: Initial total population size
# Each simulation will start with a vector of equal number of individuals in each
    stage, i.e. N0/k

popsim <- function(matarray, v, N0=100){
  k <- dim(matarray)[1]
  tmax <- dim(matarray)[3]
  Nmat<-Vmat<-matrix(NA, nrow=k, ncol=tmax)
  Nmat[,1] <- rep(N0/k,k)
  Vmat[,1] <- rep(N0/k,k)*v
  for(i in 2:tmax){
    Nmat[,i] <- matarray[,,i-1]%*%Nmat[,i-1] #population size at time i
    Vmat[,i] <- Nmat[,i]*v #total reproductive value at time i
  }
  list("N"=Nmat,"V"=Vmat)
}
```

```
********************************************************************************
Function Simulation_onepop returns the asymptotic and stochastic population growth
rates, the dominant eigenvalue of the mean matrix across environments, and
            the variance of the annual population growth rates
      - using the matrix.array and pop.sim functions defined above -
********************************************************************************
#Inputs:
#mats: a list class object which provides MatA, MatF and MatU for each matrix
    population model
#nsim: number of stochastic simulations
#tmax: number of time steps
#varZ: standard deviation (z follows a normal distribution with a mean of 0 and a
    standard deviation of 1)
#survival.link: Choice of link function for survival (logistic or loglog)
#fertility.link: Choice of link function for fertility (logistic, loglog, or log)
#betaS: Effect of z on survival on the link scale
#betaF: Effect of z on fertility on the link scale
#maxeff: Maximum fertility (if logistic or loglog)

#Outputs:
#lambda0: asymptotic population growth rate (dominant eigenvalue of the matrix
    population model)
#rS: stochastic population growth rate
#lambdaMean: dominant eigenvalue of the mean matrix across environments (estimated
    from tmax matrices)
#VarPart: the variance of the annual population growth rates, multiplied by 1/(2*
    lambda0^2)

Simulation_onepop <- function(mats, nsim=500, tmax=100, varZ=1, survival.link="
    logistic", fertility.link="logistic", betaS=surv.eff, betaF=fert.eff, maxeff= Meff
    ){
  lambda1 <- wvlambda(mats$matA)$lam
  MatA <- mats$matA/lambda1
  MatU <- mats$matU/lambda1
  MatF <- mats$matF/lambda1
  if(any(apply(MatU,2,sum)>1)){
    lambda <- wvlambda(MatA=MatA)$lam
    returns <- list("rS"= NA, "NonlinearityPart"= NA, "VariancePart"= NA, "lambda0"=
        lambda,"lambdaMean"=NA, "GenTime"=NA, "lambda0.notscaled"=lambda1)
```

```
  }
  else{
    k <- dim(MatA)[1]
    Gtime <- GenTime(MatA, MatF)
    MatR<-MatRfunc(MatU=MatU)
    MatQ<-MatQfunc(MatF=MatF)
    res <- wvlambda(MatA=MatA)
    lambda<-res$lambda
    v <- res$v #Reproductive values
    sim.mat <- matrix(NA, nrow= nsim, ncol= tmax)
    sim.mat.average <- array(dim=c(k,k,nsim))
    for(i in 1:nsim){
      zvec <- rnorm(tmax,0,sqrt(varZ))
      matarray <- matrix.array(zvec=zvec, Svec=apply(MatU,2,sum), Fvec=apply(MatF,2,
          sum) , Rmat=MatR, Qmat=MatQ, survival.link = survival.link, fertility.link =
          fertility.link, betaF=betaF,betaS=betaS, maxeff=maxeff)
      sim1 <- popsim(matarray=matarray, v=v)
      sim.mat[i,] <- apply(sim1$V,2,sum) #Total reproductive values for tmax time
          steps
      sim.mat.average[,,i]<- apply(matarray, c(1,2), mean) #average matrix across
          environments for simulation i
    }
    diffmat <- apply(log(sim.mat),1,diff) #matrix with nsim columns and tmax-1 growth
        increments of the total reproductive values for each simulation (rows)
    sim.mat.averageEnv <- apply(sim.mat.average, c(1,2), mean) #average matrix across
        nsim simulations
    lambdaMean <- wvlambda(MatA=sim.mat.averageEnv)$lambda #lambda of the average
        matrix across nsim simulations
    varDiffmean <- mean(apply(diffmat,2,var))

    returns <- list("rS"=mean(apply(diffmat,2,mean)), "NonlinearityPart"=log(
        lambdaMean)-log(lambda), "VariancePart"=1/(2*lambda^2)*varDiffmean, "lambda0"=
        lambda, "lambda0.notscaled"=lambda1, "lambdaMean"=lambdaMean, "GenTime"=Gtime)
      }
  return(returns)
}
```

```
*******************************************************************
#Function return.simulations applies the function Simulation_onepop to
#each of the models in a comadre subset.
*******************************************************************
#Inputs:
#comadremodels: an object with a subset of entries from the comadre database, with the
     same structure.
#varZ: Variance of z.
#survival.link: Choice of link function for survival (logistic or loglog)
#fertility.link: Choice of link function for fertility (logistic, loglog, or log)
#betaS: Effect of z on survival on the link scale
#betaF: Effect of z on fertility on the link scale
#maxeff: Maximum fertility (if logistic or loglog)

return.simulations <- function(comadremodels=xdataALL, varZ=1, survival.link="logistic
    ", fertility.link="logistic", betaS=surv.eff, betaF=fert.eff, maxeff= Meff, nsim
    =500, tmax=100){
  len <- length(comadremodels$mat)
  output <- data.frame(rS = rep(NA, len),
                    NonlinearityPart = rep(NA, len),
                    VariancePart = rep(NA, len),
                    lambda0 = rep(NA, len),
                    lambdaMean = rep(NA, len),
                    GenTime = rep(NA, len),
                    lambda0.notscaled = rep(NA, len))
  for (i in 1:length(comadremodels$mat)){
    mats <-comadremodels$mat[[i]]
    print(i)
    if (any(is.na(mats$matA))|any(is.na(mats$matU))|any(is.na(mats$matF))){
      output$rS[i] <- NA
      output$NonlinearityPart[i] <- NA
      output$VariancePart[i] <- NA
      output$lambda0[i] <- NA
      output$lambdaMean[i] <- NA
      output$GenTime[i] <- NA
      output$lambda0.notscaled[i] <-NA
    }
    else{
```

```
      simulres <- Simulation_onepop(mats, nsim= nsim, varZ=varZ, tmax= tmax, survival.
          link = survival.link, fertility.link = fertility.link, betaS=betaS, betaF=
          betaF, maxeff=maxeff)
      output$rS[i] <- simulres$rS
      output$NonlinearityPart[i] <- simulres$NonlinearityPart
      output$VariancePart[i] <- simulres$VariancePart
      output$lambda0[i] <- simulres$lambda0
      output$lambdaMean[i] <- simulres$lambdaMean
      output$GenTime[i] <- simulres$GenTime
      output$lambda0.notscaled[i] <- simulres$lambda0.notscaled
    }
  }
  output
}
```

```
================================================================================
                                    EXAMPLE
================================================================================

*****************************************************************************
#Step 1 - Selection of the 154 matrix population models from COMADRE database
#(cf Supporting information 1)
*****************************************************************************
#Set the working directory and load the comadre database
setwd("/...YourPath/NameOfYourFolder")
load("COMADRE_v.4.20.5.0.RData")

#Subsetting function (from Salguero-Gomez et al. 2016)
subsetDB <- function(db=comadre,sub=1:100){
  subsetID <- sub
  ssdb <- db
  ssdb$metadata <- ssdb$metadata[subsetID,]
  ssdb$mat <- ssdb$mat[subsetID]
  ssdb$matrixClass <- ssdb$matrixClass[subsetID]
  return(ssdb)
}

ListOfMatrixID <- c(240307, 240308, 240309, 240310, 240646, 240647, 240648, 240651,
    249118, 249119, 249122, 249133, 249159, 249160, 249182, 249191, 249214, 249248,
    249249, 249250, 249251, 249272, 249274, 249286, 249292, 249332, 249376, 249410,
    249411, 249412, 249464, 249504, 249506, 249512, 249524, 249525, 249605, 249619,
    249620, 249621, 249625, 249674, 249734, 249757, 249758, 249759, 249810, 249811,
    249812, 249837, 249851, 249871, 249876, 249877, 249880, 249881, 249882, 249897,
    249909, 249921, 249922, 248391, 248392, 248398, 248404, 248405, 248406, 248413,
    248433, 248441, 248446, 248451, 248452, 248454, 248473, 248478, 248515, 248518,
    248552, 248553, 248575, 248615, 248616, 248631, 248635, 248688, 248696, 248700,
    248702, 248715, 248718, 248722, 248763, 248770, 248787, 248818, 249020, 249049,
    249052, 250003, 250023, 250027, 250029, 250037, 250041, 250047, 250057, 250058,
    250060, 250088, 250094, 250114, 250115, 250116, 250118, 250120, 250122, 250123,
    250127, 250128, 250129, 250130, 250131, 250132, 250133, 250134, 250158, 250163,
    250164, 250168, 248236, 248240, 248046, 248048, 248057, 248091, 248104, 248109,
    248111, 248128, 248150, 248151, 248152, 248154, 248155, 248156, 248166, 248169,
    248173, 248178, 248186, 248220, 248221, 248222)
```

```
data <- comadre$metadata[comadre$metadata$MatrixID %in% ListOfMatrixID,]
iddata <- as.numeric(rownames(data))
#Subset the COMADRE database to select the 154 matrix population models only:
ComadreMPM154 <- subsetDB(comadre,iddata)
length(c(table(ComadreMPM154$metadata$CommonName))) #107 different species
length(c(table(ComadreMPM154$metadata$MatrixID))) #154 matrices

#Some adjustments
#Golden ground squirrel
j <- which(ComadreMPM154$metadata$CommonName == "Golden-mantled ground squirrel")
ComadreMPM154$mat[[j]]$matA[1,1] <- ComadreMPM154$mat[[j]]$matF[1,1]<- 0.2325
ComadreMPM154$mat[[j]]$matA[1,2:6] <- ComadreMPM154$mat[[j]]$matF[1, 2:6] <- 1.014927

#Great northern diver
which(ComadreMPM154$metadata$CommonName== "Great northern diver")
ComadreMPM154$mat[[19]]$matF[1,1] <- 0
ComadreMPM154$mat[[19]]$matU[1,1] <- 0.57

ComadreMPM154$mat[[20]]$matF[1,1] <- 0
ComadreMPM154$mat[[20]]$matU[1,1] <- 0.57

#Green-rumped parrotlets
ComadreMPM154$mat[[17]]$matA[1,1:2] <- c(0.451,0.612)
ComadreMPM154$mat[[17]]$matA[2,1:2] <- c(0.121,0.915)
ComadreMPM154$mat[[17]]$matF[1,1:2] <- c(0,0.501)
ComadreMPM154$mat[[17]]$matF[2,1:2] <- c(0,0.1357)
ComadreMPM154$mat[[17]]$matU[1,1:2] <- c(0.451,0.111)
ComadreMPM154$mat[[17]]$matU[2,1:2] <- c(0.121,0.7793)

# Exclude bony fish species from the selection (N=132 matrices)
ComadreMPM154_noFish <- ComadreMPM154
ComadreMPM154_noFish$metadata <- ComadreMPM154$metadata[-c(which(ComadreMPM154$
    metadata$Class == "Actinopterygii")),]
ComadreMPM154_noFish$matrixClass <- ComadreMPM154$matrixClass[-c(which(ComadreMPM154$
    metadata$Class == "Actinopterygii"))]
ComadreMPM154_noFish$mat <- ComadreMPM154$mat[-c(which(ComadreMPM154$metadata$Class ==
     "Actinopterygii"))]
```

```
******************************************************************
#Step 2 - ANALYTICAL APPROACH
#Return all the stochastic demographic entities (no bony fish MPMs)
******************************************************************
#Positive covariance, betaS = betaF = 0.4
#varZ = 1
Meff <- 2.5

AA_Logis_beta0.4_var1_Posi <- return.results(comadremodels= ComadreMPM154_noFish,
varZ= 1, survival.link= "logistic", fertility.link= "logistic", betaS= 0.4,
betaF= 0.4, maxeff= Meff)

AA_Logis_beta0.4_var1_Posi <- data.frame(Species = ComadreMPM154_noFish$metadata$
    CommonName, AA_Logis_beta0.4_var1_Posi)

AA_Logis_beta0.4_var1_Posi <- AA_Logis_beta0.4_var1_Posi[complete.cases(AA_Logis_beta0
    .4_var1_Posi),]

*****************************************************************
#Step 3 - SIMULATION APPROACH
#Return all the stochastic demographic entities (no bony fish MPMs)
******************************************************************

#Stochastic perturbations using S and F rates as logistic functions of z
#nsim=100 (instead of 4000), tmax=200
#Positive covariance, betaS = betaF = 0.4
#Meff = 2.5 and varZ = 1

SA_Logis_beta0.4_var1_Posi <- return.simulations(comadremodels=ComadreMPM154_noFish,
varZ= 1, survival.link= "logistic", fertility.link= "logistic", betaS= 0.4,
betaF= 0.4, maxeff= Meff, nsim= 100, tmax= 200)

SA_Logis_beta0.4_var1_Posi <- data.frame(Species = ComadreMPM154_noFish$metadata$
    CommonName, SA_Logis_beta0.4_var1_Posi)

SA_Logis_beta0.4_var1_Posi <- SA_Logis_beta0.4_var1_Posi[complete.cases(SA_Logis_beta0
    .4_var1_Posi),]
```

```
========
Plot
========
library(ggthemes)
library(ggplot2)
library(reshape2)

## e.g., ANALYTICAL APPROACH

#Plot 1 - Effects of increased environmental variability on the stochastic population
    growth rate across generation time (GenTime), when (st)age-specific survival
    probabilities were logistic functions of the environment z.

ggplot(AA_Logis_beta0.4_var1_Posi, aes(x =GenTime, y = rS)) + geom_point(size=1)+
    geom_segment(aes(x = 0, y = 0, xend = 60, yend = 0),
                col=8, lwd=0.5, linetype="dashed")+
    theme_tufte(base_family ='')+
    geom_rangeframe(data = data.frame(GenTime = c(0, 60), rS = c(-0.1, 0.05))) +
    scale_y_continuous(breaks = c(-0.1,-0.05,0,0.05), limits = c(-0.1, 0.05))+
    scale_x_continuous(breaks = c(0,20,40,60), limits = c(-0.01, 62))+
    theme( plot.title = element_text(face = "bold", size = rel(1.81), hjust = 0.5),
            axis.title = element_text(face = "bold",size = rel(1.3)),
            axis.text = element_text(size = rel(1.2)),
            axis.title.y = element_text(),
            axis.title.x = element_text(),
            plot.margin=unit(c(3,6,3,3),"mm"))+
    guides(color = guide_legend(override.aes = list(size = 2)))+
    labs(y=expression(paste("ln(",lambda["s"], ")")), x="Generation time")
```

```
#Plot 2 - Decomposition of the stochastic growth rate into main components capturing
    variance-covariance and nonlinearity effects on ln(lambda_bar)

*************
AA_Logis_beta0.4_var1_Posi.melt <- AA_Logis_beta0.4_var1_Posi[,c(2,9,10,5)]

AA_Logis_beta0.4_var1_Posi.melt$VariancePart <- AA_Logis_beta0.4_var1_Posi.melt$
    VariancePart*-1
names(AA_Logis_beta0.4_var1_Posi.melt)[2:4] <- c("Nonlinearity in fj(z)","Nonlinearity
     in sj(z)", "Variance-covariance [fj(z)+sj(z)]")
AA_Logis_beta0.4_var1_Posi.melt1 <- melt(AA_Logis_beta0.4_var1_Posi.melt, id.vars= c('
    GenTime'), measure.vars= c("Nonlinearity in sj(z)","Nonlinearity in fj(z)", "
    Variance-covariance [fj(z)+sj(z)]"), variable.name= 'Part', value.name= 'Contri')
*************

ggplot(AA_Logis_beta0.4_var1_Posi.melt1, aes(x =GenTime, y = Contri)) + geom_point(
    size=1, aes(col=Part, shape=Part))+
  geom_segment(aes(x = 0, y = 0, xend = 60, yend = 0),
               col=8, lwd=0.5, linetype="dashed")+
  theme_tufte(base_family ='')+
  scale_colour_manual(values=c("darkgoldenrod1","red", "blue")) + scale_shape_manual(
      values=c(19,19, 17)) +
  geom_rangeframe(data = data.frame(GenTime = c(0, 60), Contri = c(-0.125, 0.10))) +
  scale_y_continuous(breaks = c(-0.10,0,0.10), limits = c(-0.125, 0.10))+
  scale_x_continuous(breaks = c(0,20,40,60), limits = c(-0.01, 62))+
  theme( plot.title = element_text(face = "bold", size = rel(1.81), hjust = 0.5),
        axis.title = element_text(face = "bold",size = rel(1.3)),
        axis.text = element_text(size = rel(1.2)),
        axis.title.y = element_text(vjust = 0.5,size = rel(0.9)),
        axis.title.x = element_text(),
        legend.title = element_blank(),
        legend.text = element_text(size = rel(1.2)),
        legend.key.width = unit(0.3,"cm"),
        legend.key.size = unit(0.3, "cm"),
        legend.position = c(0.72,0.075),
        plot.margin=unit(c(3,3,3,3),"mm"))+
  guides(color = guide_legend(override.aes = list(size = 2)))+
  labs(y=expression(paste("Contribution to ln(",lambda["s"], ")")),
      x="Generation time")
```
